# Deep-learning-based cell composition analysis from tissue expression profiles

**DOI:** 10.1101/659227

**Authors:** Kevin Menden, Mohamed Marouf, Sergio Oller, Anupriya Dalmia, Karin Kloiber, Peter Heutink, Stefan Bonn

## Abstract

We present Scaden, a deep neural network for cell deconvolution that uses gene expression information to infer the cellular composition of tissues. Scaden is trained on single cell RNA-seq data to engineer discriminative features that confer robustness to bias and noise, making complex data preprocessing and feature selection unnecessary. We demonstrate that Scaden outperforms existing deconvolution algorithms in both precision and robustness. A single trained network reliably deconvolves bulk RNA-seq and microarray, human and mouse tissue expression data and leverages the combined information of multiple data sets. Due to this stability and flexibility, we surmise that deep learning will become an algorithmic mainstay for cell deconvolution of various data types. Scaden’s comprehensive software package is easy to use on novel as well as diverse existing expression datasets available in public resources, deepening the molecular and cellular understanding of developmental and disease processes.

## Introduction

The analysis of tissue-specific gene expression using Next Generation Sequencing (RNA-seq) is a centerpiece of the molecular characterization of biological and medical processes (*1*). A well-known limitation of tissue-based RNA-seq is that it typically measures average gene expression across many molecularly diverse cell types that can have distinct cellular states (*2*). A change in gene expression between two conditions can therefore be attributed to a change in the cellular composition of the tissue or a change in gene expression in a specific cell population, or a mixture of the two. To deconvolve the cell type composition from a change in gene expression is especially important in systems with cellular proliferation (e.g. cancer) or cellular death (e.g. neuronal loss in Neurodegenerative Diseases) due to systematic cell population differences between experimental groups (*3*).

To account for this problem, several computational cell deconvolution methods have been proposed during the last years (*4, 5*). These algorithms utilize gene expression profiles (GEPs) of cell type-specifically expressed genes to estimate cellular fractions using linear regression in order to detect, interpret, and possibly correct for systematic differences in cellular abundance between samples (*4*). While the best performing linear regression algorithms for deconvolution seem to be variations of Support Vector Regression (SVR) (*6–10*), the selection of an optimal GEP is a field of active research (*10, 11*). Indeed, it has been recently shown that the design of the GEP is the most important factor in most deconvolution methods, as results from different algorithms strongly correlate given the same GEP (*11*).

In theory, an optimal GEP should contain a set of genes that are predominantly expressed within each cell population of a complex sample (*12*). They should be stably expressed across experimental conditions, for example across health and disease, and resilient to experimental noise and bias. However, bias is typically inherent to biomedical data and is imparted, for instance, by intersubject variability, variations across species, different data acquisition methods, different experimenters, or different data types. The negative impact of bias on deconvolution performance can be partly improved by using large, heterogeneous GEP matrices (*11*). It is therefore not surprising that recent advancement in cell deconvolution relied almost exclusively on sophisticated algorithms to normalize the data and engineer optimal GEPs (*10*).

While GEP-based approaches lay the foundational basis of modern cell deconvolution algorithms, we hypothesize that Deep Neural Networks (DNNs) could create optimal features for cell deconvolution, without relying on the complex generation of GEPs. DNNs such as multilayer perceptrons are universal function approximators that achieve state-of-the-art performance on classification and regression tasks. Whereas this feature is of little importance for strictly linear input data, it makes DNNs superior to linear regression algorithms as soon as data deviates from ideal linearity. This means, for instance, that as soon as data is noisy or biased and classical linear regression algorithms may falter, the hidden layer nodes of the DNN learn to represent higher-order latent representations of cell types that do not depend on input noise and bias. We theorize, therefore, that by using gene expression information as network input, hidden layer nodes of the DNN would represent higher-order latent representations of cell types that are robust to input noise and technical bias.

An obvious limitation of DNNs is the requirement for large training data to avoid overfitting of the machine learning model. While ground truth information on tissue RNA-seq cell composition is scarce, one can use single cell RNA-seq (scRNA-seq) data to obtain virtually unlimited *in silico* tissue datasets of predefined cell composition (*7–9, 13–15*). We do this by sub-sampling and subsequently merging cells from scRNA-seq datasets, this approach being limited only by the availability of tissue-specific scRNA-seq data. It is to be noted that scRNA-seq data suffers from biases, such as drop-out, that RNA-seq data is not subject to(*16*). While this complicates the use of scRNA-seq data for GEP design (*8*), we surmise that latent network nodes could represent features that are robust to such biases.

Based on these assumptions we developed a single-cell-assisted deconvolutional DNN (Scaden) that uses simulated bulk RNA-seq samples for training and predicts cell type proportions for input expression samples of cell mixtures. Scaden is trained on publicly available scRNA- and RNA-seq data, does not rely on specific GEP matrices, and automatically infers informative features. Finally, we show that Scaden deconvolves expression data into cell types with higher precision and robustness than existing methods that rely on GEP matrices.

## Results

### Scaden Overview, Model Selection, and Training

In this part we focus on the design and optimization of Scaden by training, validation, and testing on *in silico* data. Note that the generation of in silico data is a strictly linear mathematical operation. Our aim in this context, in order to corroborate Scaden’s basic functionality, is to show that Scaden’s performance compares with (but not necessarily exceeds) that of state-of-the-art algorithms.

The basic architecture of Scaden is a DNN that takes gene counts of RNA-seq data as input and outputs predicted cell fractions (Fig. 1). To optimize the performance of the DNN, it is trained on data that contains both the gene expression and the real cell type fraction information (Fig. 1B). The network then adjusts its weights to minimize the error between the predicted cell fractions and the real cell fractions (Fig. 1C).

**Figure 1.**
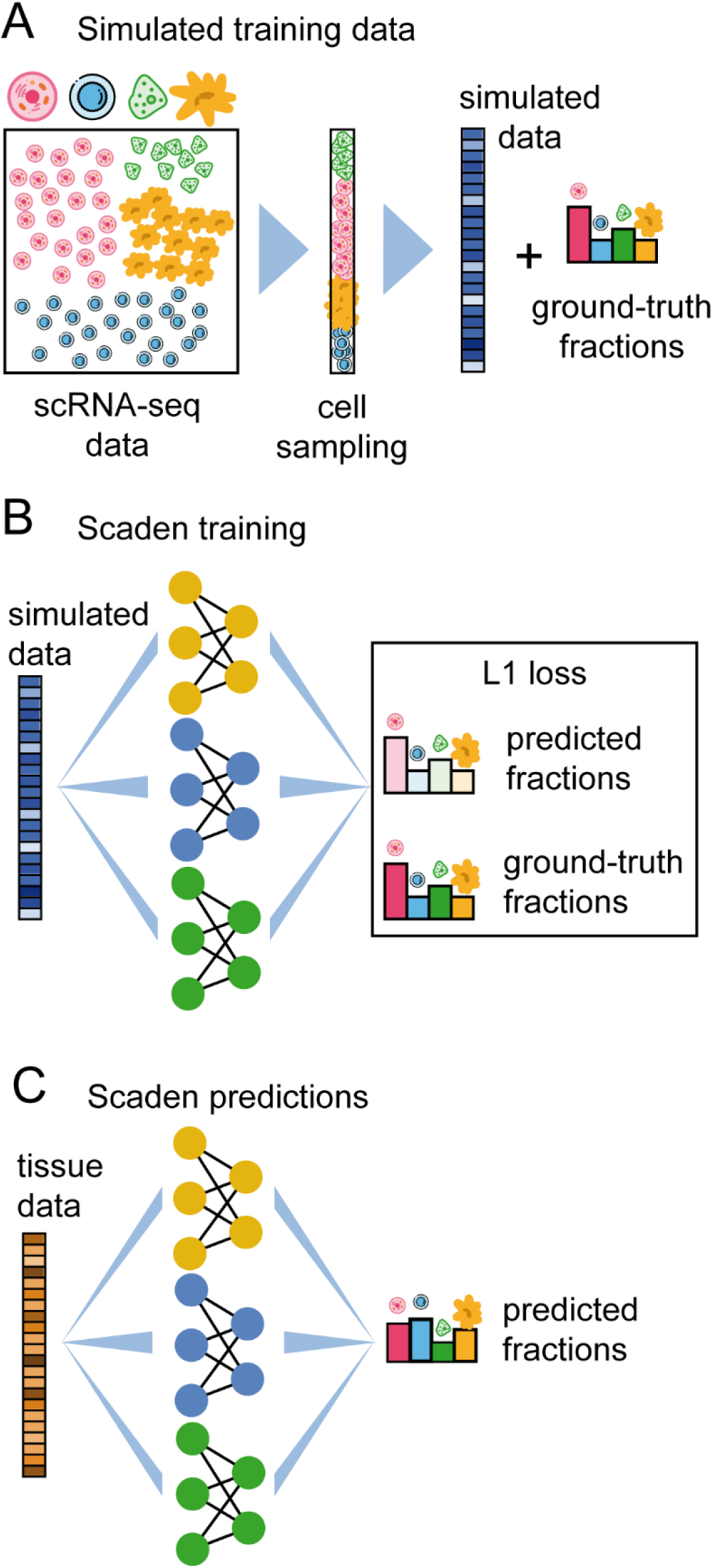
Overview of training data generation and cell type deconvolution with Scaden. A: Artificial bulk samples are generated by subsampling random cells from a scRNA-seq datasets and merging their expression profiles. B: Model training and parameter optimization on simulated tissue RNA-seq data by comparing cell fraction predictions to ground-truth cell composition. C: Cell deconvolution of real tissue RNA-seq data using Scaden.

For the model selection and training we made use of the virtually unlimited amount of artificial bulk RNA-seq datasets with defined composition that can be generated *in silico* from published scRNA-seq and RNA-seq datasets (simulated tissues) (Fig. 1A, Tables S1 & S2). The only constraint is that the scRNA-seq and RNA-seq data must come from the same tissue as the bulk data subject to deconvolution.

To find the optimal DNN architecture for cell deconvolution, we generated bulk PBMC (peripheral blood mononuclear cells) RNA-seq data from four publicly available scRNA-seq data sets (Tables S1 & S3). We performed leave-one-dataset-out cross validation, training Scaden on mixtures of synthetic datasets from three scRNA-seq datasets and evaluating the performance on simulated tissue from a fourth scRNA-seq dataset.

We used the root mean square error (RMSE), Pearson’s correlation coefficient (r), the slope and intercept of the regression fitted for ground truth and predicted cell fractions, and Lin’s concordance correlation coefficient (CCC) (*17*) to assess algorithmic performance. The CCC is a measure sensitive not only to scatter but also to deviations from linearity (slope and intercept). Within the main text, we report on CCC and RMSE values only, other metrics can be found in the supplements.

The final Scaden model is an ensemble of the three best performing models (Table S4) and the final cell type composition estimates are the averaged predictions of all three ensemble models (Fig. 1 & S1). Using an ensemble of models increased the deconvolution performance as compared to single best models (Table S6). Details of the model and hyper-parameters are given in Table S5. We also evaluated the effect of the size of the training data set on Scaden deconvolution performance, repeating leave-one-dataset-out cross validation on PBMC data with training dataset sizes from 150 up to 15,000 samples (Fig. S2). The increase in CCC value starts to level off from about 1,500 simulated samples for this data set but continues to increase slowly with sample size. We specifically addressed the question to what degree the DNN, trained on simulated samples, tends to overfit, failing to generalize to real bulk RNA-seq data. To understand after how many steps a model trained on *in silico* data overfits on real RNA-seq data, we trained Scaden on simulated data from an ascites scRNA-seq dataset (Table S1, 6,000 samples) and evaluated the loss function on a corresponding annotated RNA-seq dataset (*18*) (Table S2, 3 samples) as a function of the number of steps (Fig. S3). All models converged after approximately 5,000 steps, and slightly overfit when trained for longer. Based on this result, we opted for an early-stop approach after 5,000 steps for evaluation on real bulk RNA-seq data.

We then compared Scaden to four state-of-the-art GEP-based cell deconvolution algorithms, CIBERSORT (CS) (*6*), CIBERSORTx (CSx) (*7*), MuSiC (*8*), and Cell Population Mapping (CPM) (*9*). While CS relies on hand-curated GEP matrices, CSx, MuSiC, and CPM can generate GEPs using scRNA-seq data as input.

To get an initial estimate of Scaden’s deconvolution fidelity we trained the model on 24,000 simulated PBMC RNA-seq samples from three datasets and tested its performance in comparison to CS, CSx, MuSiC and CPM on a fourth dataset of 500 samples each (e.g. training on data6k, data8k, donorA and evaluation on donorC). We used corresponding scRNA-seq data sets for the construction of GEPs as input for CSx and MuSiC, and CPM. For CS we used the PBMC-optimized LM22 GEP matrix(*6*), which was developed by the CS authors for the deconvolution of human PBMC data.

For two of four test datasets (donorA, donorC), Scaden obtained the highest CCC and lowest RMSE, followed by CSx, MuSiC, CS, and CPM (Fig. S4, Table S7). CSx and MuSiC obtained the highest CCC values for the data8k and data6k datasets, respectively. Scaden obtained the highest average CCC and lowest RMSE (0.88, 0.08, respectively), followed by MuSiC (0.85, 0.10), CSx (0.83, 0.11), CS (0.63 0.15), and CPM (0, 0.20) (Fig. S4). As expected, all algorithms that use scRNA-seq data as reference performed well, with the notable exception of CPM. We want to mention that CPM focuses on the reconstruction of continuous spectra of cellular states, while it incorporates cell deconvolution as an additional feature. We therefore report CPM’s deconvolution performance in the supplementary material from here on. On average, Scaden also obtained the highest correlation and the best intercept and slope values on simulated PBMC data (Table S7). A closer inspection on a per cell type bases (Fig. 2A) revealed that Scaden yields consistently higher CCC values and lower RMSEs when compared to the other algorithms.

**Figure 2.**
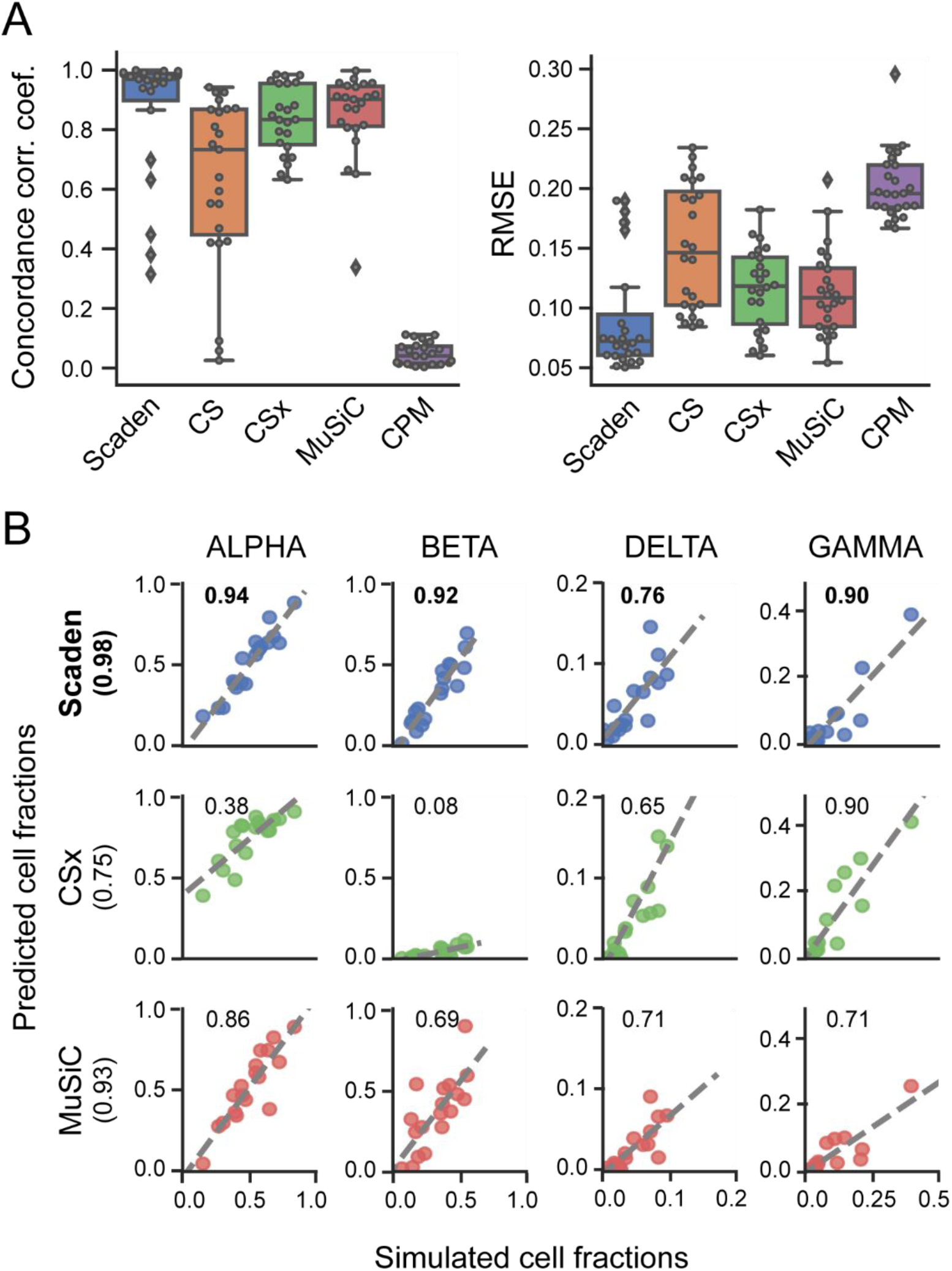
Deconvolution performance on simulated tissue data. A: Boxplots of the cell type prediction concordance correlation coefficient (CCC) and RMSE for four simulated PBMC datasets. Tables S14 and S16 contain information on the five (six for CS) cell types used. B: Scatterplots for four pancreas cell types of ground truth (x-axis) and predicted values (y-axis) for Scaden, CSx, and MuSiC on artificial pancreas data(*19*). Numbers inside the plotting area signify CCC values.

A specific feature of the MuSiC algorithm is that it preferentially weighs genes according to low inter-subject and intra-cell cluster variability for its GEP, which increases deconvolution robustness when high expression heterogeneity is observed between human subjects, for example (*8*). To understand if Scaden can utilize multi-subject information to increase its deconvolution performance, we trained Scaden, CSx, and MuSiC on scRNA-seq pancreas data from several subjects (*20*) and assessed the performance on a separate simulated pancreas RNA-seq dataset (*19*). To allow for direct comparison, we chose the same pancreas training and test datasets that were used in the original MuSiC publication (Table S1). To enable Scaden to leverage the heterogeneity of multi-subject data, training data was generated separately for every subject in the dataset (see Methods). CSx cannot profit from multi-subject data but performed well on the artificial PBMC datasets and was therefore included in the comparison. The best average performance (across cell types) is achieved by Scaden (CCC = 0.98), closely followed by MuSiC (CCC = 0.93), while CSx does not perform as well (CCC = 0.75) (Fig. 2B, Table S8). On a per cell-type basis, Scaden’s predictions are clearly superior to the other two algorithms for all cell types. This provides strong evidence that Scaden, by separating training data generation for each subject, can learn inter-subject heterogeneity and outperform specialized multi-subject algorithms such as MuSiC on the cell-type deconvolution task.

Additionally, we wanted to test how the best performing deconvolution algorithms Scaden, MuSiC, and CSx behave when unknown cell content is part of the mixture. To test this, all cells falling into the ‘Unknown’ category were removed from the training or reference PBMC datasets but added to the simulated mixture samples at fixed percentages (5%, 10%, 20%, 30%) (see Methods). Scaden obtains the highest CCC for all tested percentages of unknown cell content (Fig. S5, Table S9). The general deconvolution performance declines linearly with increasing percentage of unknown content for all tested algorithms, indicating that Scaden, MuSiC, and CSx have a similar robustness against unknown mixture content.

### Robust deconvolution of bulk expression data

The true use case of cell deconvolution algorithms is the cell fraction estimation of tissue RNA-seq data. Especially for noisy and biased bulk RNA-seq data we hypothesize that Scaden’s latent feature representations might help it to more robustly predict cell fractions as compared to GEP-based algorithms.

We therefore assessed the performance of Scaden, CS, CSx, and MuSiC to deconvolve two publicly available human PBMC bulk RNA-seq datasets, for which curated GEP matrices as well as RNA-seq data with associated ground truth cell type compositions from flow cytometry are available. We will refer to these datasets that consists of 12 samples each as PBMC1 (*21*) and PBMC2 (*10*) (Table S2). Deconvolution for all methods was performed as described in the previous section, with the difference that data from all four PBMC scRNA-seq datasets was now deployed for Scaden training. Results are given in Fig. 3A, B & C and Tables S10 & S11.

**Figure 3.**
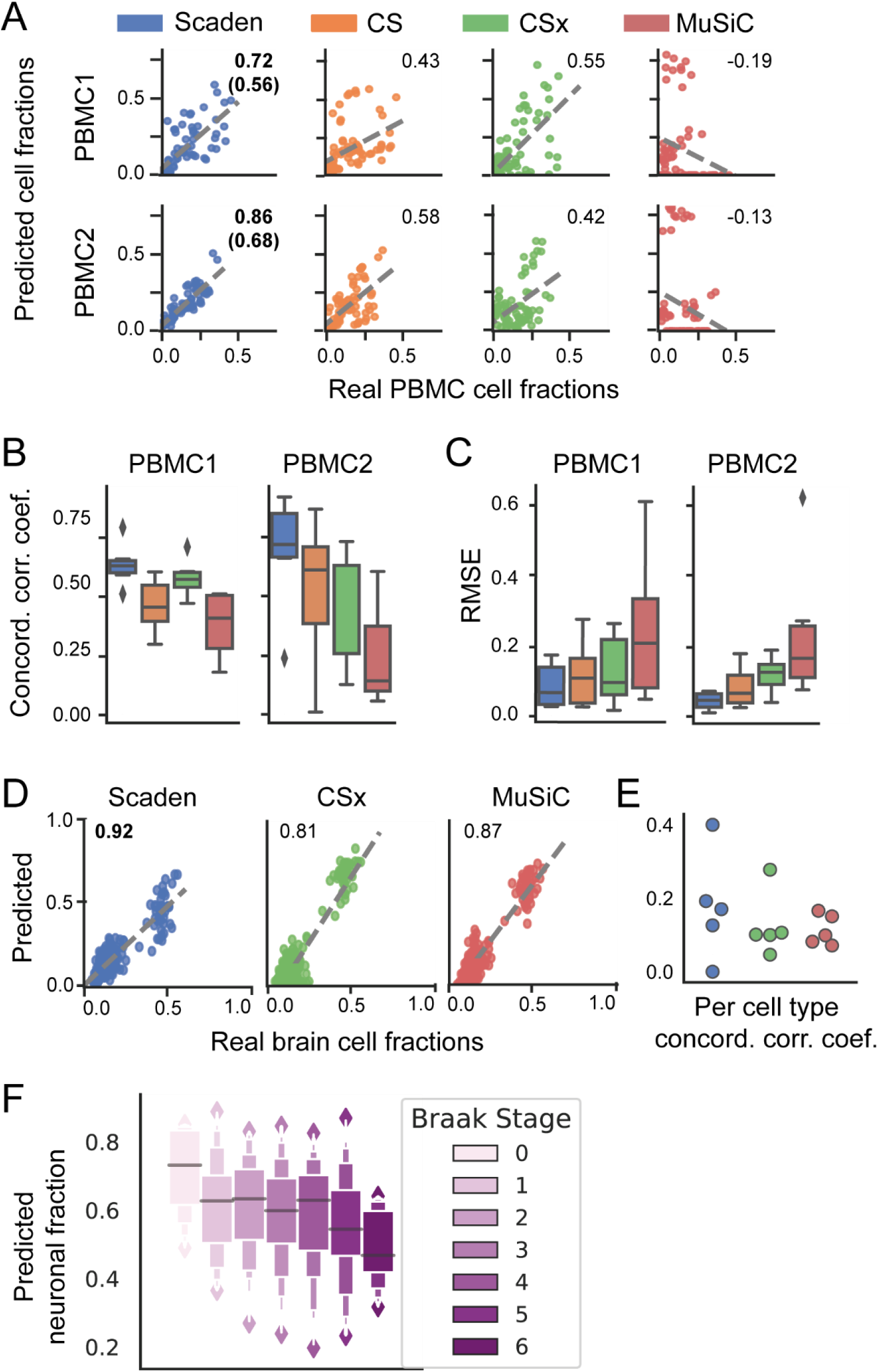
Deconvolution of real tissue RNA-seq data. A: Per-cell-type scatterplots of ground truth (x-axis) and predicted values (y-axis) for Scaden, CS, CSx and MuSiC on real PBMC1 and PBMC2 cell fractions. Numbers inside the plotting area signify CCC values. For Scaden, the CCC using only scRNA-seq training data (in parenthesis) and the CCC using mixed scRNA-seq and RNA-seq training data is shown. B: Boxplots of CCC values for real PBMC1 (first column) and PBMC2 (second column) data. C: RMSE values for real PBMC1 (first column) and PBMC2 (second column) data. D: Prediction of human brain cell fractions of the ROSMAP dataset using the Darmanis data set as reference: Scatterplots of ground truth (x-axis) and predicted values (y-axis) for Scaden, CSx, MuSiC, of data. CCC values are shown as inserts. E: Per cell type CCC values for ROSMAP using the Darmanis data as reference. F: Neuronal content determined by Scaden trained on mouse brain data and evaluated on the Braak stage of the ROSMAP study.

On the PBMC1 dataset and using all cell types, Scaden obtained the highest CCC and lowest RMSE (0.56, 0.13), while CSx (0.55, 0.16) and CS (0.43, 0.15) performed well yet significantly worse than Scaden (Fig. 3A, Tables S10 & S11). CPM (0, 0.18) and MuSiC (−0.19, 0.32) both failed to deconvolve the cell fractions of the PBMC1 data. Scaden also obtained the best CCC and RMSE (0.68, 0.08) on the PBMC2 dataset, while CS (0.58, 0.10) and CSx (0.42, 0.13) obtained good deconvolution results. Similar to the PBMC1 data deconvolution results, CPM (−0.16, 0.11) as well as MuSiC (−0.13, 0.30) did not perform well on the PBMC2 deconvolution task. In addition to CCC and RMSE metrics, Scaden achieves the best correlation, intercept and slope on both PBMC datasets (Tables S10 & S11).

In particular, Scaden outperforms classical algorithms on a per cell-type basis (Fig. 3B & C). These results show weaker correlations and a strong dependence on the cell type. A closer examination of the metrics in Table S11 and Figure S6 shows that the largest variations are found in the slope and intercept.

An additional algorithmic feature of Scaden is that it seamlessly integrates increasing amounts of training data, which can be of different types, such as a combination of simulated tissue and real tissue data with cell fraction information. In theory, even limited real tissue training data could make Scaden robust to data type bias and consequently improve Scaden’s deconvolution performance on real tissue data. We therefore trained Scaden on a mix of simulated PBMC and real PBMC2 (12 samples) data and evaluated its performance on real PBMC1 data (Fig. 3A & B, S6, Tables 10 & S11). While the training contained very little (∼2%) real data, Scaden’s CCC increased from 0.56 to 0.72 and the RMSE decreased from 0.13 to 0.10. We observed similar performance increases when Scaden was trained on simulated PBMC and real PBMC1 data and evaluated on real PBMC2 data (Fig. 3A & B, S6, Tables 10 & S11). This further validates that Scaden reliably deconvolves tissue RNA-seq data into the constituent cell fractions and that very accurate deconvolution results can be obtained if reference and target datasets are from the same experiment.

We next wanted to test how the algorithm performs on post-mortem human brain tissue of a subsample from the ROSMAP study (*22*), for which ground truth cell composition information was recently measured by immunohistochemistry (41 samples with all cell types given) (*23*). The data provided by this study consists of bulk RNA-seq data from the dorsolateral prefrontal cortex (DLPFC) and poses a special challenge due to the complexity of its cell type composition, which is further complicated by the fact that data originates from brains of healthy individuals as well as AD (Alzheimer Disease) patients at various stages of neuronal loss. As reference datasets, we used the scRNA-seq dataset provided by Darmanis *et al*. (*24*) from the anterior temporal lobe of living patients and the Lake dataset that isolates nuclei of neurons from two (visual and frontal) cortical regions from a postmortem brain and subjects them to RNA sequencing (*25*). From these, we generated 2000 training samples (Darmanis) and 4000 samples (two regions from the Lake dataset).

Fig. 3D shows the deconvolution results for all three algorithms with the Darmanis (scRNA-seq) reference dataset. Scaden achieves the highest CCC value (0.92) followed by MuSiC (0.87) and CSx (0.81) (Table S12). Compared to Scaden, MuSiC and CSx overestimate neural percentages, leading to higher RMSE values of 0.09 and 0.12, respectively (Scaden: 0.06). Notably, all methods showed a lower concordance correlation coefficient on the per cell type level (Fig. 3E), demonstrating that some per-cell-type correlations are poor, either in slope, intercept, variance, or a combination of them. This emphasizes the need for a cell type-specific inspection of results and highlights that, depending on the dataset, cell type-specific deconvolution results can be far from perfect.

In addition to comparing the predictive power of Scaden, CSx and MuSiC on human brain tissue with different reference datasets, we also tested how the choice of reference datasets affected Scaden’s deconvolution results. Notably, all methods significantly drop in performance when the Lake snRNA-seq dataset is used as reference as we had presumed (Figure S7A). We want to emphasize that Scaden, in contrast to CSx and MuSiC, has the possibility to simultaneously use both datasets as reference, whereas for CSx and MuSiC, the user has to choose one of the two, unaware which will give the correct results.

Indeed, we found that the performance of Scaden was almost unaffected when the Lake dataset was added to the training samples (CCC=0.90, RMSE=0.06) (Figure S7A, Table S12). Finally, when calculating the CCC values on a per-sample basis, Scaden achieves the best scores for most samples (Figure S7B).

In a next step, we wanted to assess whether Scaden’s deconvolution performance was robust across species by trying to predict the cell fractions of the ROSMAP study (*22*) with a Scaden model trained on *in silico* data from five mouse brain scRNA-seq datasets (Table S1). Intriguingly, Scaden was able to achieve a CCC value of 0.83 and an RMSE of 0.079 (Figure 3D, left panel), showing that Scaden can reliably deconvolve RNA-seq data across related species.

The ROSMAP study also contains information on the Braak stages (*26*) corresponding to 390 human post-mortem prefrontal cortex samples, which correlate with the severity and progression stage of AD and the degree of neuronal loss. We used the Scaden model trained on artificial data generated from five mouse brain scRNA-seq datasets to predict neuronal cell fractions of this larger human dataset. Overall, Scaden’s cell fraction predictions capture the increased neuronal loss with increasing Braak stage (Fig. 3F). Interestingly, the largest drop in neural percentage is observed at stage 5, when the neurodegeneration typically reaches the prefrontal cortex of the brain.

Given the robustness with which Scaden predicts tissue RNA-seq cell fractions using scRNA-seq training data, even across species, we next wanted to investigate if a scRNA-seq-trained Scaden model can also deconvolve other data types. To this end, we measured the deconvolution performance on a bulk PBMC microarray dataset (20 samples)(*6*) of a Scaden model trained on scRNA-seq and RNA-seq PBMC data (see above). We compared Scaden to CS using the microarray-derived LM22 matrix. CS achieved a slightly higher CCC and slightly lower total RMSE (0.72, 0.11) than Scaden (0.71, 0.13), while Scaden obtained the highest average CCC (0.50) compared to CS (0.39) (Fig. S8, Table S13). Notably in this scenario, Scaden was trained entirely on simulated scRNA-seq and RNA-seq data, while CS’s LM22 GEP was optimized on PBMC microarray data.

Overall, we provide strong evidence that Scaden robustly deconvolves tissue data across tissues, species, and even data types.

## Discussion

Scaden is a novel deep learning-based cell deconvolution algorithm that in many instances compares favorably in both prediction robustness and accuracy to existing deconvolution algorithms that rely on GEP design and linear regression. We believe that Scaden’s performance relies to a large degree on the inherent feature engineering of the DNN. The network does not only select features (genes) for regression, it also creates novel features that are optimal for the regression task in the nodes of the hidden layers. These hidden features are non-linear combinations of the input features (gene expression), which makes it notoriously difficult to explain how a DNN works (*27*). It is important to highlight that this feature creation is fundamentally different from all other existing cell deconvolution algorithms, which rely on heuristics that select a defined subset of genes as features for linear regression.

Another advantage of this inherent feature engineering is that Scaden can be trained to be robust to input noise and bias (e.g. batch effects). Noise and bias are all prevalent in experimental data, due to different sample quality, sample processing, experimenters, and instrumentation, for example. If the network is trained on different datasets of the same tissue, however, it learns to create hidden features that are robust to noise and bias, such as batch effects. This robustness is pivotal in real world cell deconvolution use cases, where the bulk RNA data for deconvolution and the training data (and therefore the network and GEP) contain different noise and biases. While especially recent cell deconvolution algorithms include batch correction heuristics prior to GEP construction, Scaden optimizes its hidden features automatically when trained on data from various batches.

The robustness to noise and bias, which might be due to hidden feature generation, is especially evident in Scaden’s ability to deconvolve across data types. A network trained on *in silico* bulk RNA-seq data can seamlessly deconvolve microarray data of the same tissue. This is quite noteworthy, as microarray data is known to have a reduced dynamic range and several hybridization-based biases compared to RNA-seq data. In other words, Scaden can deconvolve bulk data of types it has never been trained on, even in the face of strong data type bias. This raises the possibility that Scaden trained on scRNA-seq data might reliably deconvolve other bulk omics data as well, such as proteomic and metabolomic data. This assumption is strengthened by the fact that Scaden, trained on scRNA-seq data, attains state-of-the-art performance on the deconvolution of bulk RNA-seq data, two data types with very distinct biases (*16*).

As highlighted in the introduction, a drawback for many DNNs is the large amount of training data required to obtain robust performance. Here, we used scRNA-seq data to create virtually unlimited amounts of *in silico* bulk RNA-seq data of predefined type (target tissue) with known composition, across datasets. This immediately highlights Scaden’s biggest limitation, the dependency on scRNA-seq data of the target tissue. In this study we have shown that Scaden, trained solely on simulated data from scRNA-seq datasets, can outperform GEP-based deconvolution algorithms. We did observe, however, that the addition of labeled RNA-seq samples to the training data did significantly improve deconvolution performance in the case of PBMC data. We therefore believe that efforts to increase the similarity between simulated training data and the target bulk RNA-seq data could increase Scaden’s performance further. Mixtures of *in silico* bulk RNA-seq data and publically available RNA-seq data, of purified cell types for example, could further increase the deconvolution performance of Scaden. Furthermore, domain adaptation methods can be used to improve performance of models that are trained on data (here, scRNA-seq data) that is similar to the target data (here, RNA-seq data) (*28*). In future versions, Scaden’s simple multilayer perceptron architecture could leverage domain adaptation to further stabilize and improve its cell deconvolution performance.

Recent cell deconvolution algorithms have used cell fraction estimates to infer cell type-specific gene expression from bulk RNA-seq data. It is straightforward to use Scaden’s cell fraction estimates to infer per group (*3*) and per sample (*7*) cell type-specific gene expression using simple regression or non-negative matrix factorization, respectively. We would like to add a note of caution, however, as the error of cell fraction estimates, which can be quite significant, is propagated into the gene expression calculations and will affect any downstream statistical analysis.

In summary, the deconvolution performance, robustness to noise and bias, the flexibility to learn from large numbers of *in silico* datasets, across data types (scRNA-seq and RNA-seq mixtures), and potentially even tissues makes us believe that DNN-based architectures will become an algorithmic mainstay of cell type deconvolution.

## Methods

### Datasets and pre-processing

#### scRNA-seq datasets

The following human PBMC scRNA-seq datasets were downloaded from the 10X Genomics data download page: 6k PBMCs from a Healthy Donor, 8k PBMCs from a Healthy Donor, Frozen PBMCs (Donor A), Frozen PBMCs (Donor C) (*29*). Throughout this paper, these datasets are referred to with the handles data6k, data8k, donorA and donorC, respectively. It was not intended to incorporate as many datasets as possible. Instead, these four datasets were chosen with the goal to dispose of a set of samples with consistent cell types and gene expression. This limited our choice to datasets that displayed clearly identifiable cell types for the majority of cells. The Ascites scRNA-seq dataset was downloaded from https://figshare.com as provided by Schelker(*18*). Pancreas and mouse brain datasets were downloaded from the scRNA-seq dataset collection of the Hemberg lab (https://hemberg-lab.github.io/scRNA.seq.datasets/). The human brain datasets from Darmanis *et al.* and Lake *et al.* where downloaded from GEO with accession numbers GSE67835 and GSE97930, respectively. A table listing all datasets including references to the original publications can be found in Table S1.

#### scRNA-seq preprocessing and analysis

All datasets were processed using the Python package Scanpy (v. 1.2.2) (*30*) following the Scanpy’s reimplementation of the popular Seurat’s clustering workflow. First, the corresponding cell-gene matrices were filtered for cells with less than 500 detected genes, and genes expressed in less than 5 cells. The resulting count matrix for each dataset was filtered for outliers with high or low numbers of counts. Gene expression was normalized to library size using the Scanpy function ‘normalize_per_cell’. The normalized matrix of all filtered cells and genes was saved for the subsequent data generation step.

The following processing and analysis steps had the sole purpose of assigning cell type labels to every cell. All cells were clustered using the louvain clustering implementation of the Scanpy package. The louvain clustering resolution was chosen for each dataset, using the lowest possible resolution value (low resolution values lead to less clusters) for which the calculated clusters separated the cell types appropriately. The top 1000 highly variable genes were used for clustering, which were calculated using Scanpy’s ‘filter_genes_dispersion’ function with parameters min_mean=0.0125, max_mean=3 and min_disp=0.5. Principal Component Analysis (PCA) was used for dimensionality reduction.

To identify cell types, marker genes were investigated for all cell types in question. For PBMC datasets, useful marker genes were adopted from public resources such as the Seurat tutorial for 2700 PBMCs(*31*). Briefly, IL7R was taken as marker for CD4 T-cells, LYZ for Monocytes, MS4A1 for B-cells, GNLY for Natural Killer cells, FCER1A for Dendritic cells and CD8A and CCL5 as markers for CD8 T-cells. For all other scRNA-seq datasets, marker genes and expected cell types were inferred from the original publication of the dataset. For instance, to annotate cell types of the mouse brain dataset from Zeisel *et al*. (*32*), we used the same marker genes as Zeisel and colleagues. We did not use the same cell type labels from the original publications because a main objective was to assure that cell type labeling is consistent between all datasets of a certain tissue.

Cell type annotation was performed manually across all the clusters for each dataset, such that all cells belonging to the same cluster were labeled with the same cell type. The cell type identity of each cluster was chosen by crossing the cluster’s highly differentially expressed genes with the curated cell type’s marker genes. Clusters that could not be clearly identified with a cell type were grouped into the ‘Unknown’ category.

#### Tissue Datasets for Benchmarking

To assess the deconvolution performance on real tissue expression data, we used datasets for which the corresponding cell fractions were measured and published. The first dataset is the PBMC1 dataset which was obtained from Zimmermann *et al.*(*21*). The second dataset, PBMC2, was downloaded from GEO with accession code GSE107011(*10*). This dataset contains both RNA-seq profiles of immune cells (S4 cohort) and from bulk individuals (S13 cohort). As we were interested in the bulk profiles, we only used 12 samples from the S13 cohort from this data. Flow cytometry fractions were collected from the Monaco *et al.* publication.

In addition to the above mentioned two PBMC datasets, we used Ascites RNA-seq data. This dataset was kindly provided by the authors and cell type fractions for this dataset were taken from the supplementary materials of the publication (*18*).

For the evaluation on pancreas data, artificial bulk RNA-seq samples created from the scRNA-seq dataset of Xin *et al*. (*19*) were used. This dataset was downloaded from the resources of the MuSiC publication(*8*). The artificial bulk RNA-seq samples used for evaluation were then created using the ‘bulk_construct’ function of the MuSiC tool. To assess how Scaden and the GEP algorithms deal with the presence of unknown cell types, we generated PBMC bulk RNA samples from the four scRNA-seq datasets (6000 each). The undefined amount of unknown cells that was generated by this approach was removed to be replaced by defined amounts of 5%, 10%, 20%, 30% of unknown cells, respectively. Cell fractions of all four samples were predicted with Scaden trained on the other three.

Performance on these samples was then assessed to test robustness against unseen cell types in the bulk mixture. Scaden was trained on samples from all datasets but the test dataset, while CSx and MuSiC used data8k as a reference.

The microarray dataset GSE65133 was downloaded from GEO, and cell type fractions taken from the original CS publication (*6*).

Finally, we wanted to get insights into neurodegenerative cell fraction changes in the brain. While it is known that neurodegenerative diseases like Alzheimer’s Disease are accompanied by a gradual loss of brain neurons, stage-specific cell type shifts are still hard to come by. Here we use the ROSMAP (Religious Orders Study and Memory and Aging Project Study) cortical RNA-seq dataset along with the corresponding clinical metadata, to infer cell type composition over six clinically relevant stages of neurodegeneration (*22*). Furthermore, to assess deconvolution accuracy on post-mortem human brain tissue, we used 41 samples from the ROSMAP (Religious Orders Study and Memory and Aging Project), for which cell composition information from immunohistochemistry (*23*) was recently released and for which fractions for all cell types were reported. The ROSMAP RNA-seq data was downloaded from https://www.synapse.org/. The cell composition values were kindly provided by the authors of the study (*23*).

#### RNA-seq preprocessing and analysis

For the RNA-seq datasets analyzed in this study, we did not apply any additional processing steps, but used the obtained count or expression tables directly as downloaded for all dataset except the ROSMAP dataset. For the latter, we generated count tables from raw FastQ-files using Salmon (*33*) and the GRCh38 reference genome. FastQ-files from the ROSMAP study were downloaded from Synapse (www.synapse.org).

### Simulation of bulk RNA-seq samples from scRNA-seq data

Scaden’s deep neural network requires large amounts of training RNA-seq samples with known cell fractions. This explains why the generation of artificial bulk RNA-seq data is one of the key elements of the Scaden workflow.

In order to generate the training data, preprocessed scRNA-seq datasets were used (see section ‘Data Collection and Processing’), comprising the gene expression matrix and the cell type labels. Artificial RNA-seq samples were simulated by sub-sampling cells from individual scRNA-seq datasets - cells from different datasets were not merged into samples to preserve within-subject relationships. Datasets generated from multiple subjects were split according to subject and each sub-sampling was constrained to cells from one subject in order to capture the cross-subject heterogeneity and keep subject-specific gene dependencies.

The exact sub-sampling procedure is described in the following. First, for every simulated sample, random fractions were created for all different cell types within each scRNA-seq dataset using the random module of the Python package NumPy. Briefly, a random number was chosen from a uniform distribution between 0 and 1 using the NumPy function ‘random.rand()’ for each cell type, and then this number was divided by the sum of all random numbers created to ensure the constraint of all fractions adding up to 1:

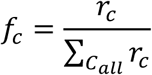

where *r_c_* is the random number created for cell type *c*, and *C_all_* is the set of all cell types. Here, *f_c_* is the calculated random fraction for cell type *c*. Then, each fraction was multiplied with the total number of cells selected for each sample, yielding the number of cells to choose for a specific cell type:

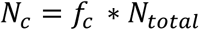

where *N_c_* is the number of cells to select for the cell type *c*, and *N_total_* is the total number of cells contributing to one simulated RNA-seq sample (500, in this study). Next, *N_c_* cells were randomly sampled from the scRNA-seq gene expression matrix for each cell type *c*. Afterwards, the randomly selected single-cell expression profiles for every cell type are then aggregated by summing their expression values, to yield the artificial bulk expression profile for this sample.

Using the above described approach, cell compositions that are strongly biased toward a certain cell type or are missing specific cell types are rare among the generated training samples. To account for this and to simulate cell compositions with a heavy bias to and the absence of certain cell types, a variation of the sub-sampling procedure was used to generate samples with sparse compositions, which we refer to as sparse samples. Before generating the random fractions for all cell types, a random number of cell types was selected to be absent from the sample, with the requirement of at least one cell type constituting the sample. After these leave-out cell types were chosen, random fractions were created and samples generated as described above. Using this procedure, we generated 32,000 samples for the human PBMC training dataset, 14,000 samples for the human pancreas training dataset, 6000 samples for human brain, and 30,000 samples for the mouse brain training dataset (Table S3).

Artificial bulk RNA-seq datasets were stored in ‘h5ad’ format using the Anndata package(*30*), which allows to store the samples together with their corresponding cell type ratios, while also keeping information about the scRNA-seq dataset of origin for each sample. This allowed to access samples from specific datasets, which is useful for cross validation.

### Scaden Overview

The following section contains an overview of the input data preprocessing, the Scaden model, model selection, and how Scaden predictions are generated.

#### Input Data Preprocessing

The data preprocessing step is aimed to make the input data more suitable for machine learning algorithms. To achieve this, an optimal preprocessing procedure should transform any input data from the simulated samples or from the bulk RNA-seq to the same feature scale. Before any scaling procedure can be applied, it must be ensured that both the training data and the bulk RNA-seq data subject to prediction share the same features. Therefore, before scaling, both datasets are limited to contain features (genes) that are available in both datasets. The two-step processing procedure used for Scaden is described in the following:

First, to account for heteroscedasticity, a feature inherent to RNA-seq data, the data was transformed into logarithmic space by adding a pseudocount of 1 and then taking the Logarithm (base 2).

Second, every sample was scaled to the range [0,1] using the MinMaxScaler() class from the Sklearn preprocessing module. Per sample scaling, unlike per feature scaling that is more common in machine learning, assures that inter-gene relative expression patterns in every sample are preserved. This is important, as our hypothesis was that a neural network could learn the deconvolution from these inter-gene expression patterns.

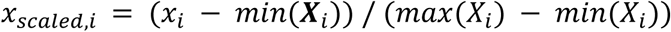

where *x_scaled,i_* is the log2 expression value of gene x in sample i, *X_i_* is the vector of log2 expression values for all genes of sample i, *min* (***X_i_***) is the minimum gene expression of vector *X_i_*, and *max* (*X_i_*) the maximum gene expression of vector *X_i_* Note that all training datasets are stored as expression values and are only processed as described above. In the deployment use-case the simulated training data should contain the same features as in the bulk RNA-seq sample that shall be deconvolved.

#### Model Selection

The goal of model selection was to find an architecture and hyperparameters that robustly deconvolve simulated tissue RNA-seq data and, more importantly, real bulk RNA-seq data. Due to the very limited availability of bulk RNA-seq datasets with known cell fractions, model selection was mainly optimized on the simulated PBMC datasets. To capture inter-experimental variation, we used leave-one-dataset-out cross validation for model optimization: a model was trained on simulated data from all but one dataset, and performance was tested on simulated samples from the left-out dataset. This allows to simulate batch effects between datasets and helps to test the generalizability of the model. In the process of model selection and (hyper-) parameter optimization, performed on PBMC and Ascites datasets, we found three models with different architectures and dropout rates but comparable performance. In order to address overfitting in individual models, we decided to use a combination of models, expecting this to serve as another means of regularization. We did not test multiple combinations, but rather used an informed choice with varying layer sizes and dropout regularization, with the goal to increase model diversity. We observed that the average of an ensemble of models generalized better to the test sets than individual models. Model training and prediction is done separately for each model, with the prediction averaging step combining all model predictions (Fig. S1, Tables S4 & S6). We provide a list of all tested parameters in the supplementary materials (Table S5).

#### Final Scaden Model

The Scaden model learns cell type deconvolution through supervised training on datasets of simulated bulk RNA-seq samples simulated with scRNA-seq data. To account for model biases and to improve performance, Scaden consists of an ensemble of three deep neural networks with varying architectures and degrees of dropout regularization. All models of the ensemble use four layers of varying sizes between 32 and 1024 nodes, with dropout-regularization implemented in two of the three ensemble models. The exact layer sizes and dropout rates are listed in Table S4. The Rectified Linear Unit (ReLU) is used as activation function in every internal layer. We used a Softmax function to predict cell fractions, as we did not see any improvements in using a linear output function with consecutive non-negativity correction and sum-to-one scaling. Python (v. 3.6.6) and the TensorFlow library (v. 1.10.0) were used for implementation of Scaden. A complete list of all software used for the implementation of Scaden is provided in Table S15.

#### Training and Prediction

After the preprocessing of the data a Scaden ensemble can be trained on simulated tissue RNA-seq data or mixtures of simulated and real tissue RNA-seq data. Parameters are optimized using Adam with a learning rate of 0.0001 and a batch size of 128. We used an L1 loss as optimization objective:

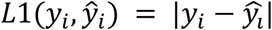

where *y_i_* is the vector of ground truth fractions of sample *i* and 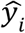 is the vector of predicted fractions of sample *i*. Each of the three ensemble models is trained independently for 5,000 steps. This ‘early stopping’ serves to avoid domain overfitting on the simulated tissue data, which would decrease the model performance on the real tissue RNA-seq data. We observed that training for more steps lead to an average performance decrease on real tissue RNA-seq data. To perform deconvolution with Scaden, a bulk RNA-seq sample is fed into a trained Scaden ensemble and three independent predictions for the cell type fractions of this sample are generated by the trained deep neural networks. These three predictions are then averaged per cell type to yield the final cell type composition for the input bulk RNA-seq sample:

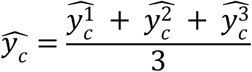

where 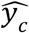 is the final predicted fraction for cell type *c* and 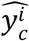 is the predicted fraction for cell type *c* of model *i*.

#### Scaden requirements

Currently, a disadvantage of the Scaden algorithm is the necessity to train a new model for deconvolution if no perfect overlap in the feature space exists. This constraint limits the usefulness of pre-trained models. Once trained, however, the prediction runtime scales linearly with sample numbers and is usually in the order of seconds, making Scaden a useful tool if deconvolution is to be performed on very large datasets. While the requirements are dataset dependent, the Scaden demo was profiled to require a peak of 3.2GB of RAM during the DNN training process, so a computer with 8GB of RAM should be able to run it smoothly. In our tests with an Intel(R) Xeon(R) CPU E5-1630 workstation the demo could run in 22 minutes, spending most of the CPU time in the DNN training process. The most prominent and obvious issue of Scaden is the difference between simulated scRNA-seq data used for training and the bulk RNA-seq data subject to inference. While Scaden is able to transfer the learned deconvolution between the two data types and achieves state-of-the-art performance, we hypothesize that efforts to improve this translatability could improve Scaden’s prediction accuracy even further. Algorithmic improvements are therefore likely to address this issue and are planned for future releases

### Algorithm Comparison

We used several performance measures to compare Scaden to four existing cell deconvolution algorithms, CIBERSORT with LM22 GEP (CS), CIBERSORTx (CSx), MuSiC and CPM. To compare the performance of the five deconvolution algorithms we measured the root mean squared error (RMSE), Lin’s concordance correlation coefficient *CCC*, Pearson product moment correlation coefficient *r*, and *R*^2^ values comparing real and predicted cell fractions estimates. Additionally, to identify systematic prediction errors and biases, slope and intercept for the regression lines were calculated. These metrics are defined as follows:

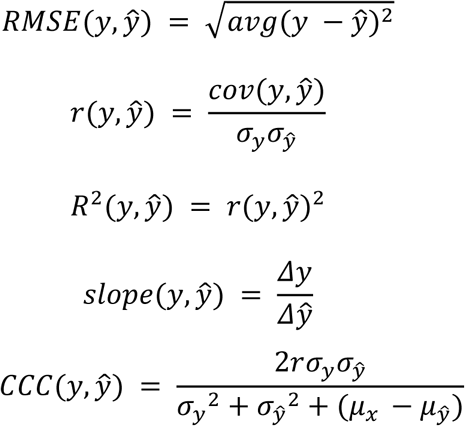

where *y* are the ground truth fractions, 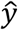 are the prediction fractions, *σ_x_* is the standard deviation of 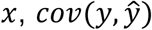 is the covariance of *y* and 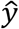, and 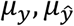 are the mean of the predicted and ground truth fractions, respectively.

All metrics were calculated for all data points of a dataset, and separately for all data points of a specific cell type. For the latter approach, we then averaged the resulting values to recover single values. While the metrics calculated on all data points might be sufficient, we deem that the cell type-specific deconvolution might in many instances be of even greater interest. It is noteworthy in this context that cell type-specific deconvolution performance can be quite weak, depending on the dataset. This is true for all tested deconvolution algorithms, while Scaden achieves best performance.

#### CIBERSORT (CS)

CS is a cell convolution algorithm based on specialized GEPs and support vector regression. Cell composition estimations were obtained using the CS web application (https://cibersort.stanford.edu/). For all deconvolutions with CS, we used the LM22 GEP, which was generated by the CS authors from 22 leukocyte subsets profiled on the HGU133A microarray platform.

Because the LM22 GEP matrix contains cell types at a finer granularity than what was used for this study, predicted fractions of sub-cell types were added together. For cell grouping, we used the mapping of sub-cell types to broader types given by Figure 6 from Monaco *et al*. (*10*). We provide a table with the exact mappings used here in the supplementary material (Table S13). The deconvolution was performed using 500 permutations with quantile normalization disabled for all datasets but GSE65133 (Microarray), as is recommended for RNA-seq data. We used default settings for all other CS parameters.

#### CIBERSORTx (CSx)

CSx is a recent variant of CS that can generate GEP matrices from scRNA-seq data and use these for deconvolution. For additional deconvolution robustness, it applies batch normalization to the data. All signature matrices were created by uploading the labeled scRNA-seq expression matrices and using the default options. Quantile normalization was disabled. For deconvolution on simulated data, no batch normalization was used. For all bulk RNA-seq datasets, the S-Mode batch normalization was chosen. All PBMC datasets were deconvolved using a GEP matrix generated from the data6k dataset (for simulated samples from data6k, a donorA GEP matrix was chosen).

#### MuSiC

MuSiC is a deconvolution algorithm that uses multi-subject scRNA-seq datasets as GEP matrices in an attempt to include heterogeneity in the matrices to improve generalization. While MuSiC tries to address similar issues of previous deconvolution algorithms by using scRNA-seq data, the approach is very different. For deconvolution, MuSiC applies a sophisticated GEP-based deconvolution algorithm that uses weighted non-negative least squares regression with an iterative estimation procedure that imposes more weight on informative genes and less weight on non-informative genes.

The MuSiC R package contains functionality to generate the necessary GEP matrix given a scRNA-seq dataset and cell type labels. To generate MuSiC deconvolution predictions on PBMC datasets, we used the data8k scRNA-seq dataset as reference data for MuSiC and follow the tutorial provided by the authors to perform the deconvolution. For deconvolution of artificial samples generated from the data8k dataset, we provided MuSiC with the data6k dataset as reference instead.

MuSiC was developed with a focus on multi-subject scRNA-seq datasets, in which the algorithm tries to take advantage from the added heterogeneity that these datasets contain, by calculating a measure of cross-subject consistency for marker genes. To assess how Scaden performs on multi-subject datasets compared to MuSiC, we evaluated both methods on artificial bulk RNA-seq samples from human pancreas. We used the ‘bulk_construct’ function from MuSiC to combine the cells from all 18 subjects contained in the scRNA-seq dataset from Xin *et al.* to generate artificial bulk samples for evaluation. Next, as a multi-subject reference dataset, we used the pancreas scRNA-seq dataset from Segerstolpe *et al*. (*20*), which contains single-cell expression data from 10 different subjects, 4 of which with type-2 Diabetes. For Scaden, the Segerstolpe scRNA-seq dataset was split by subjects, and training datasets were generated for each subject, yielding in total 10,000 samples. For MuSiC, a processed version of this dataset was downloaded from the resources provided by the MuSiC authors(*8*) and used as input reference dataset for the MuSiC deconvolution. Deconvolution was then performed according to the MuSiC tutorial, and performance compared according to the above-defined metrics.

#### Cell Population Mapping (CPM)

CPM is a deconvolution algorithm that uses single-cell expression profiles to identify a so-called ‘cell population map’ from bulk RNA-seq data(*9*). In CPM, the cell population map is defined as composition of cells over a cell-state space, where a cell-state is defined as a current phenotype of a single cell. Contrary to other deconvolution methods, CPM tries to estimate the abundance of all cell-states and types for a given bulk mixture, instead of only deconvolving the cell types. As input, CPM requires a scRNA-seq dataset and a low-dimensional embedding of all cells in this dataset, which represents the cell-state map. As CPM estimates abundances of both cell-states and types, it can be used for cell type deconvolution by summing up all estimated fractions for all cell-states of a given cell type - a method that is implemented in the scBio R package, which contains the CPM method. To perform deconvolution with CPM, we used the data6k PBMC scRNA-seq dataset as input reference for all PBMC samples. For samples simulated from the data6k dataset, we used the data8k dataset as reference. According to the CPM paper, a dimension reduction method can be used to obtain the cell-state space. We therefore used UMAP, a dimension reduction method widely used for scRNA-seq data, to generate the cell-state space mapping for the input scRNA-seq data. Deconvolution was then performed using the CPM function of the scBio package with a scRNA-seq and accompanying UMAP embedding as input.

## Data Availability

Only publicly available datasets were used during this study. The scRNA-seq PBMC datasetse donorA, donorC, data6k and data8k were all downloaded from 10X Genomics (https://support.10xgenomics.com/single-cell-gene-expression/datasets), were they are listed as ‘Frozen PBMCs (Donor A)’, ‘Frozen PBMCs (Donor C)’, ‘6k PBMCs from a Healthy Donor’ and ‘8k PBMCs from a Healthy Donor’, respectively. The Segerstolpe *et al*. scRNA-seq pancreas dataset was downloaded from ArrayExpress with accession code E-MTAB-5061. The scRNA-seq datasets from Baron *et al*. (pancreas), Tasic *et al*., Zeisel *et al*., Romanov *et al*., Campbell *et al*. and Chen *et al*. (all mouse brain) were all downloaded from https://hemberg-lab.github.io/scRNA.seq.datasets/. The ascites scRNA-seq dataset was downloaded from https://figshare.com/s/711d3fb2bd3288c8483a. The bulk RNA-seq dataset PBMC1 is accessible from ImmPort with accession code SDY67. The PBMC2 dataset was downloaded from GEO with accession code GSE107011. The ROSMAP human brain RNA-seq dataset was downloaded from Synapse (ID: syn3219045). The bulk RNA-seq data from ascites was kindly provided by Schelker *et al*. The pancreas scRNA-seq dataset from Xin *et al*. was accessed from the MuSiC tutorial site (https://xuranw.github.io/MuSiC/articles/pages/data.html).

## Code Availability

The source code for Scaden is available at https://github.com/KevinMenden/scaden. Documentation is published at https://scaden.readthedocs.io. Code to generate the figures along with the training datasets used in this study is published at figshare: https://figshare.com/projects/Scaden/62834.

## List of abbreviations

RNA-seq: Next Generation RNA Sequencing
GEP: gene expression profile matrix
SVR: Support Vector Regression
DNN: Deep Neural Network
scRNA-seq: single cell RNA-seq
simulated tissue: training data generated by mixing proportions of scRNA-seq data
PBMC: peripheral blood mononuclear cells
CCC: concordance correlation coefficient
r: Pearson’s correlation coefficient
CS: CIBERSORT
CSx: CIBERSORTx
CPM: Cell Population Mapping

## Contributions

KM and SB initiated the project. KM, PH, and SB designed the study, deep learning models, and analysis. KM, MM, and SO built the deep learning models. KM, MM, KK, and AD analyzed the data. KM, KK, and SB wrote and MM, AD, and PH contributed to the manuscript writing.

## Acknowledgements

We would like to thank the people of the Genome Biology of Neurodegenerative Diseases group and Institute of Medical Systems Biology for helpful discussions and suggestions.

## Supplementary Figures & Tables

**Supplementary Table S1:** scRNA-seq datasets used for the generation of simulated tissues for Scaden training.

| Tissue | Name | # cells | # Subjects | Source |
| --- | --- | --- | --- | --- |
| **PBMC** | data6k | 5,419 | 1 | 10X Genomics |
| **PBMC** | data8k | 8,381 | 1 | 10X Genomics |
| **PBMC** | donorA | 2,900 | 1 | 10X Genomics |
| **PBMC** | donorC | 9,519 | 1 | 10X Genomics |
| **Mouse Brain** | Tasic | 1,679 | 1 | Tasic et al., Nat. Neurosci., 2016 |
| **Mouse Brain** | Zeisel | 3,005 | 1 | Zeisel et al., Science, 2015 |
| **Mouse Brain** | Romanov | 2,881 | 1 | Romanov et al., Nat. Neurosci., 2018 |
| **Mouse Brain** | Campbell | 21,086 | 1 | Campbell et al, Nat. Neurosci., 2017 |
| **Mouse Brain** | Chen | 14,437 | 1 | Chen et al., Cell Rep., 2017 |
| **Pancreas** | Segerstolpe | 3,514 | 10 | Segerstolpe et al., Cell Metab., 2016 |
| **Pancreas** | Baron | 8,569 | 4 | Baron et al., Cell Syst., 2016 |
| **Ascites** | Ascites | 3,114 | 3 | Schelker et al, Nat. Comm., 2018 |
| **Human Brain** | Darmanis | 465 | 1 | Darmanis et al., PNAS, 2015 |
| **Human Brain** | Lake | 27,416 | 1 | Lake et al., Science, 2016 |

**Supplementary Table S2:** Bulk tissue RNA-seq datasets used for performance evaluation

| Tissue | Name | # Samples | Reference |
| --- | --- | --- | --- |
| **PBMC** | PBMC1 | 12 | Zimmermann et al., PLOS one, 2016 |
| **PBMC** | PBMC2 | 12 | Monaco et al., Cell Reports, 2019 |
| **Pancreas** | Xin | 18 | Xin et al., Cell Metab., 2016 |
| **Human Brain** | ROSMAP | 390 | Bennett et al., Curr Alzheimer Res., 2012 |
| **Ascites** | Ascites | 3 | Schelker at al., Nat. Comm. 2018 |

**Supplementary Figure S1:**
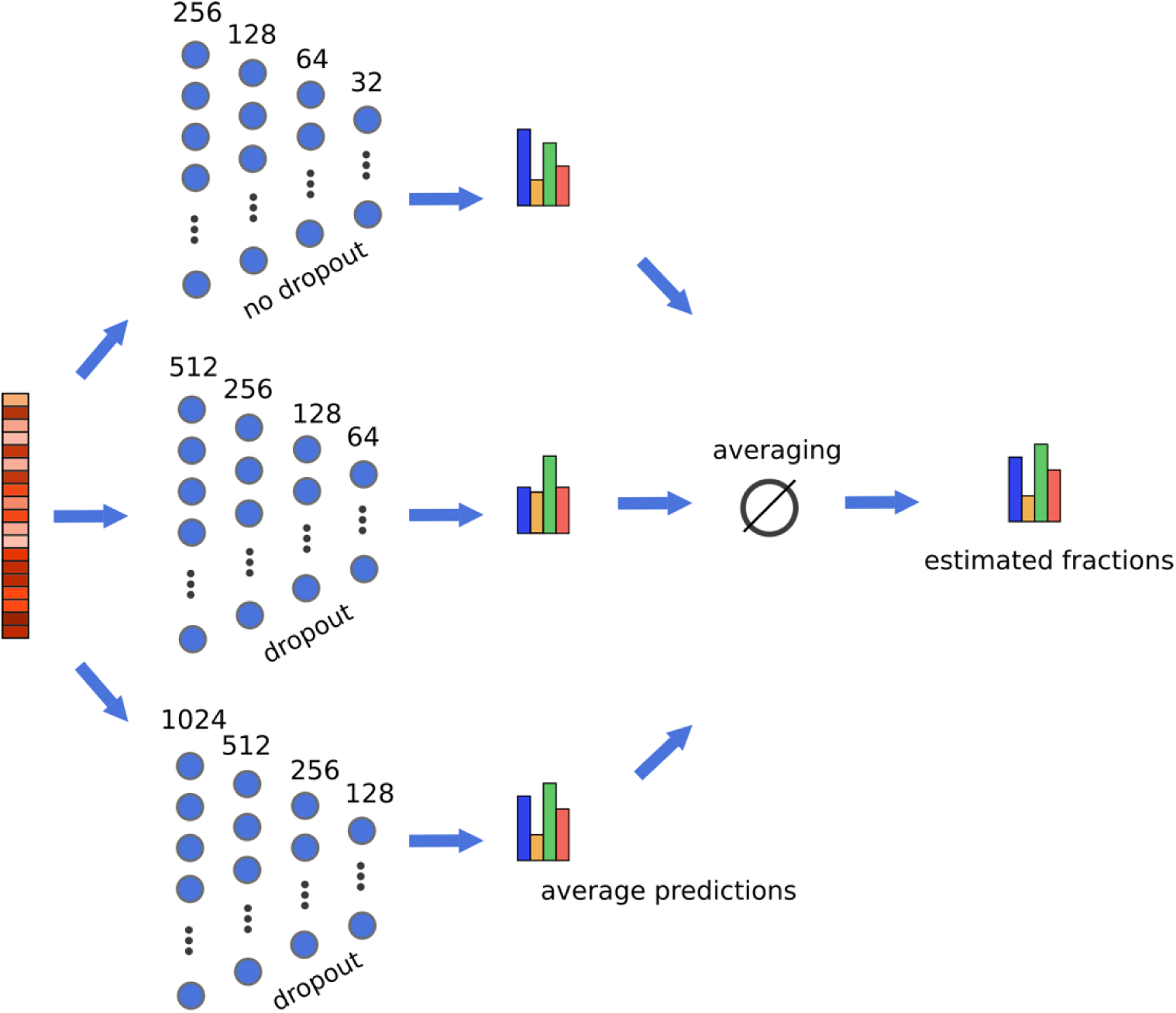
Overview of Scaden neural network ensemble model. A bulk RNA-seq sample is the input to three separate deep neural networks with varying layer sizes and dropout regularization. The predictions of all three models are subsequently averaged to obtain the final Scaden predictions. During training, predictions are not averaged and each model is trained separately.

**Supplementary Table S3:** Number of samples, datasets and size of the simulated training data.

| Tissue | # Samples | # Datasets | Size |
| --- | --- | --- | --- |
| **PBMC** | 32,000 | 4 | 1.2 GB |
| **Pancreas** | 14,000 | 2 | 0.6 GB |
| **Human Brain** | 6,000 | 2 | 0.32 GB |
| **Ascites** | 6,000 | 1 | 0.38 GB |
| **Mouse Brain** | 30,000 | 5 | 1.5 GB |

**Supplementary Table S4:** Architectures of deep neural network models used in Scaden ensemble. All models use an L1 as a loss function, ReLU activation for all layers but the last, and softmax activation for the last layer.

| Model | # Layers | Layer sizes | Dropout rates |
| --- | --- | --- | --- |
| **M256** | 4 | 256, 128, 64, 32 | 0, 0, 0, 0 |
| **M512** | 4 | 512, 256, 128, 64 | 0, 0.3, 0.2, 0.1 |
| **M1024** | 4 | 1024, 512, 256, 128 | 0, 0.6, 0.3, 0.1 |

**Supplementary Table S5:** Hyperparameters used for model optimization

| Parameter | Values tested |
| --- | --- |
| **Batch size** | 32, 64, 128, 256, 512 |
| **# Layers** | 2, 3, 4 |
| **Layer sizes** | 2048, 1024, 512, 256, 128, 64, 32, 16 |
| **Dropout rate** | [0, 0.8] |
| **Loss function** | L1, L2 |

**Supplementary Table S6:** Comparison of Scaden models and the Scaden ensemble on four PBMC scRNA-seq datasets. Concordance correlation coefficient was calculated on all datasets separately and then averaged.

|  | Scaden Ensemble | M256 | M512 | M1024 |
| --- | --- | --- | --- | --- |
| **CCC** | 0.914 | 0.898 | 0.909 | 0.907 |

**Supplementary Figure S2:**
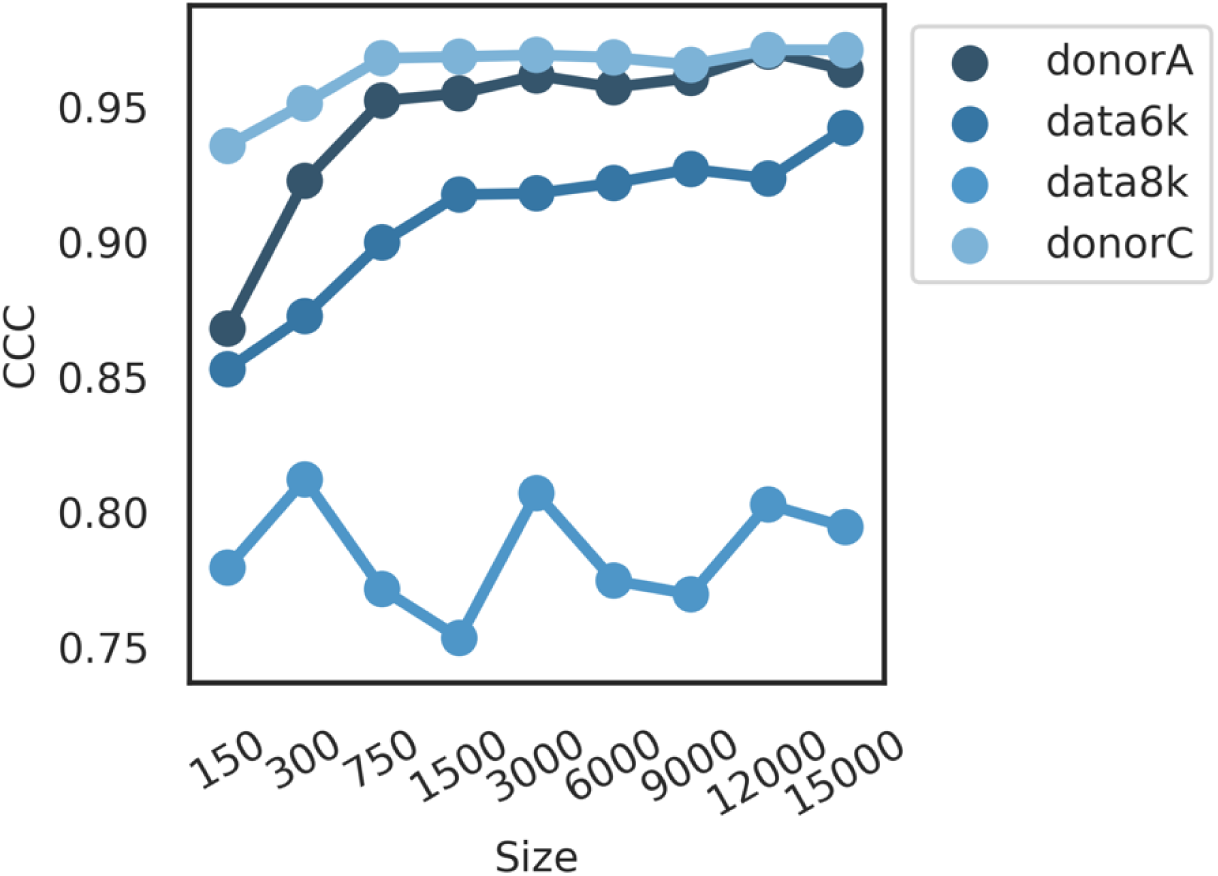
Assessment of the effect of training data size on the CCC. We used four publicly available PBMC scRNA-seq datasets (see also Table S1) to train Scaden on different sample sizes between 150 and 15,000. Each simulated RNA-seq training dataset was generated from three scRNA-seq datasets. Simulated testing datasets were generated from a separate scRNA-seq dataset.

**Supplementary Figure S3:**
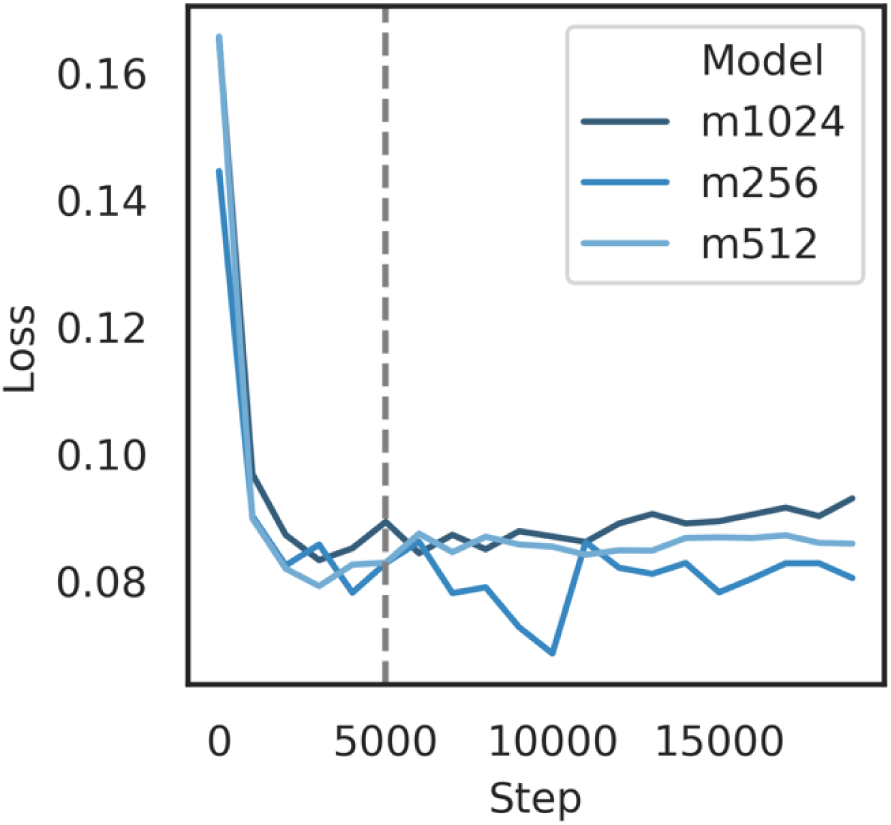
Assessment of overfitting. We trained Scaden on simulated data from ascites scRNA-seq datasets (6000 samples, see Table S1) and evaluated the loss function on annotated bulk RNA-seq datasets (see also Table S2) (3 samples) as a function of training steps. The results led us to an early-stop approach after 5,000 steps for evaluation on real bulk RNA-seq data.

**Supplementary Table S7:** Deconvolution evaluation on simulated PBMC data.

| Method | DS | RMSE | Slope | Correlation | Intercept | CCC |
| --- | --- | --- | --- | --- | --- | --- |
| CPM | data6k | 0.192 | 0.03 | 0.082 | 0.162 | 0.053 |
| CPM | data8k | 0.185 | 0.048 | 0.263 | 0.159 | 0.093 |
| CPM | donorA | 0.239 | -0.081 | -0.259 | 0.18 | -0.147 |
| CPM | donorC | 0.189 | 0.038 | 0.102 | 0.16 | 0.066 |
| CS | data6k | 0.163 | 0.508 | 0.57 | 0.082 | 0.566 |
| CS | data8k | 0.136 | 0.551 | 0.708 | 0.075 | 0.687 |
| CS | donorA | 0.137 | 0.605 | 0.767 | 0.066 | 0.746 |
| CS | donorC | 0.168 | 0.45 | 0.522 | 0.092 | 0.517 |
| CSx | data6k | 0.106 | 0.756 | 0.824 | 0.041 | 0.821 |
| CSx | data8k | 0.097 | 0.744 | 0.863 | 0.043 | 0.854 |
| CSx | donorA | 0.125 | 0.696 | 0.81 | 0.051 | 0.801 |
| CSx | donorC | 0.094 | 0.829 | 0.865 | 0.029 | 0.864 |
| MuSiC | data6k | 0.086 | 0.848 | 0.887 | 0.025 | 0.886 |
| MuSiC | data8k | 0.136 | 0.663 | 0.728 | 0.056 | 0.725 |
| MuSiC | donorA | 0.1 | 0.811 | 0.883 | 0.031 | 0.88 |
| MuSiC | donorC | 0.084 | 0.897 | 0.896 | 0.017 | 0.896 |
| Scaden | data6k | 0.104 | 0.747 | 0.83 | 0.042 | 0.825 |
| Scaden | data8k | 0.133 | 0.625 | 0.73 | 0.063 | 0.722 |
| Scaden | donorA | 0.035 | 0.92 | 0.988 | 0.013 | 0.985 |
| Scaden | donorC | 0.046 | 0.849 | 0.973 | 0.025 | 0.964 |

**Supplementary Figure S4:**
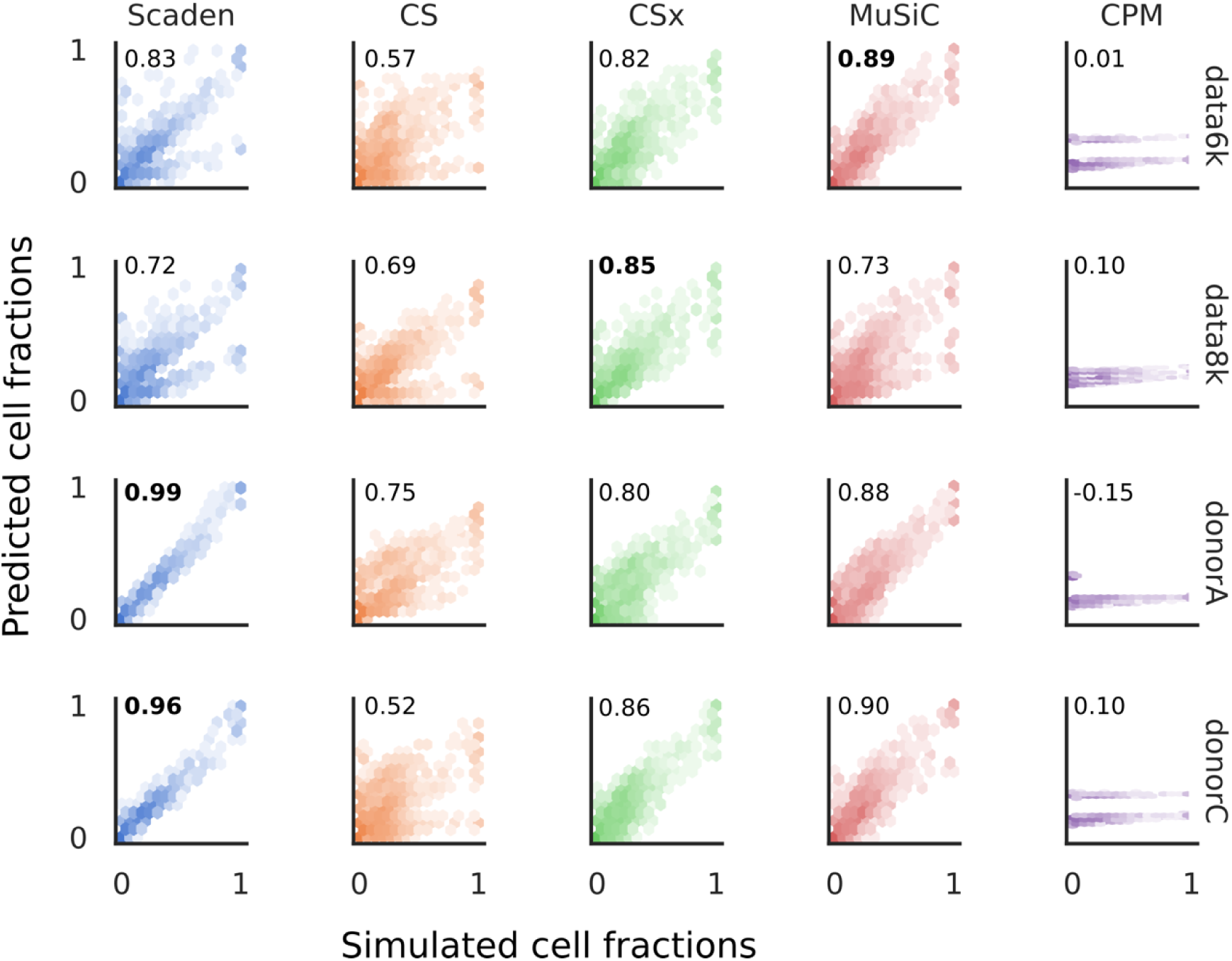
Scatter plots of predicted (y-axis) vs ground truth values (x-axis) for four simulated PBMC RNA-seq datasets (data6k, data8k, donorA, donorC) for all tested algorithms (Scden, CS, CSx, MuSiC, CPM). CCCs are given as insets in each plot.

**Supplementary Table S8:** Deconvolution performance on simulated pancreas data from Xin et al. on a per cell-type level.

| Method | Celltype | RMSE | Correlation | Slope | Intercept | CCC |
| --- | --- | --- | --- | --- | --- | --- |
| CSx | ALPHA | 0.282 | 0.816 | 0.691 | 0.431 | 0.375 |
| CSx | Average | 0.171 | 0.845 | 0.891 | 0.1 | 0.499 |
| CSx | BETA | 0.309 | 0.833 | 0.175 | -0.017 | 0.078 |
| CSx | DELTA | 0.04 | 0.812 | 1.567 | -0.013 | 0.647 |
| CSx | GAMMA | 0.052 | 0.921 | 1.131 | 0.0 | 0.897 |
| CSx | Total | 0.212 | 0.79 | 1.113 | -0.028 | 0.746 |
| MuSiC | ALPHA | 0.11 | 0.887 | 1.108 | -0.042 | 0.863 |
| MuSiC | Average | 0.087 | 0.835 | 0.861 | -0.008 | 0.744 |
| MuSiC | BETA | 0.148 | 0.752 | 1.067 | 0.017 | 0.694 |
| MuSiC | DELTA | 0.023 | 0.817 | 0.716 | -0.003 | 0.707 |
| MuSiC | GAMMA | 0.068 | 0.881 | 0.552 | -0.003 | 0.711 |
| MuSiC | Total | 0.099 | 0.938 | 1.078 | -0.019 | 0.929 |
| Scaden | ALPHA | 0.067 | 0.949 | 1.071 | -0.034 | 0.942 |
| Scaden | Average | 0.051 | 0.902 | 1.031 | -0.02 | 0.881 |
| Scaden | BETA | 0.07 | 0.936 | 1.152 | -0.045 | 0.916 |
| Scaden | DELTA | 0.024 | 0.807 | 1.012 | 0.008 | 0.764 |
| Scaden | GAMMA | 0.045 | 0.914 | 0.89 | -0.008 | 0.901 |
| Scaden | Total | 0.055 | 0.978 | 1.033 | -0.008 | 0.976 |

**Supplementary Figure S5.**
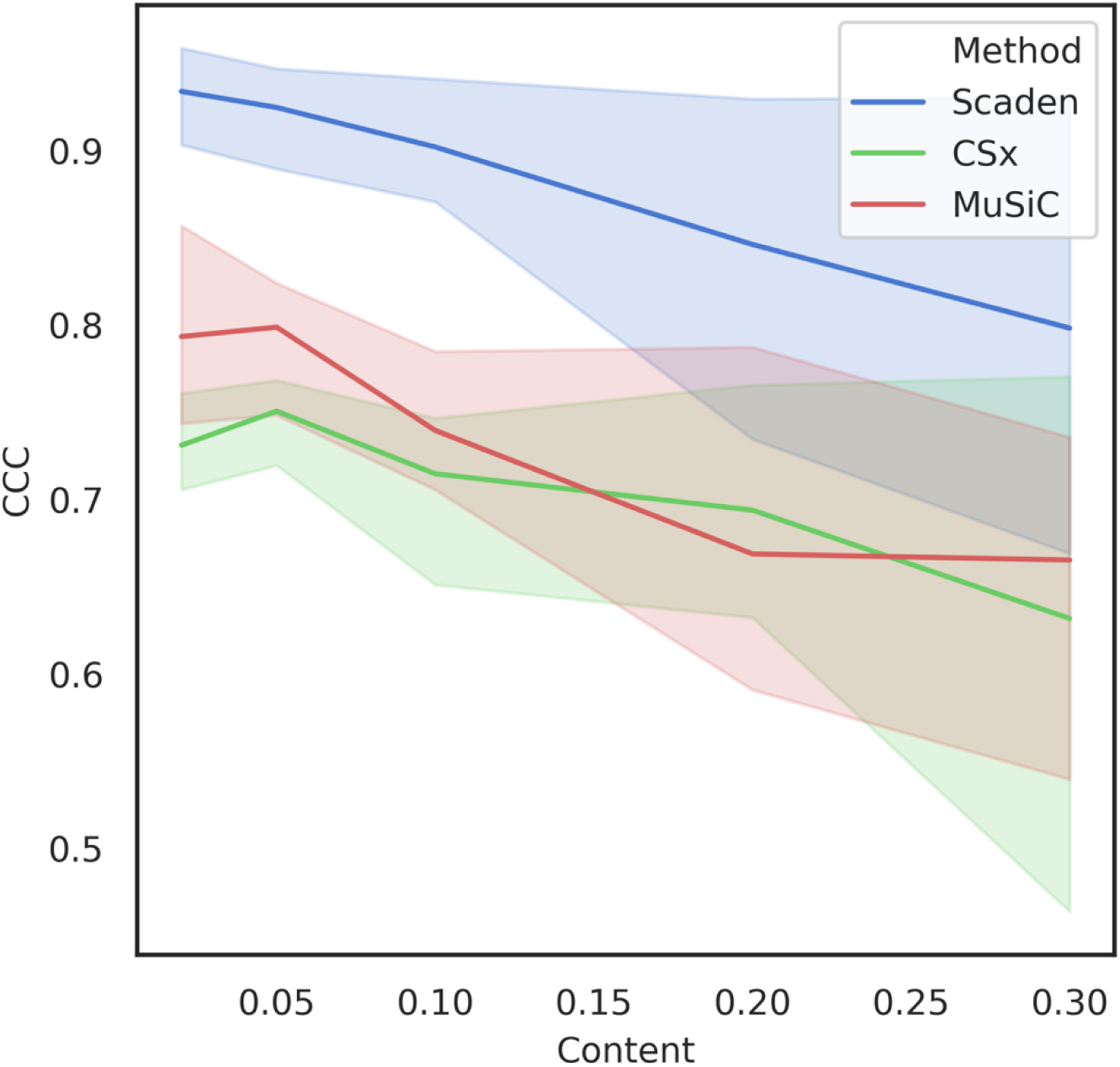
Deconvolution performance on simulated PBMC datasets with added unknown mixture contents. Cell types from 24,000 *in silico* PBMC samples (Table S1) were replaced by a defined percentage of unkown cells (5, 10, 20, 30 %), which were removed from the data prior to sample simulation. The shaded areas mark the 95% confidence interval. One dataset was evaluated by Scaden trained on *in silico* data from the remaining three. For CSx and MuSiC, a different dataset was used as reference.

**Supplementary Table S9:** Deconvolution performance on datasets with added unknown mixture contents.

| Method | Content | CCC | RMSE | Correlation | Intercept | Slope |
| --- | --- | --- | --- | --- | --- | --- |
| CSx | 0.02 | 0.731 | 0.097 | 0.738 | 0.032 | 0.841 |
| CSx | 0.05 | 0.751 | 0.092 | 0.754 | 0.035 | 0.823 |
| CSx | 0.1 | 0.715 | 0.092 | 0.719 | 0.041 | 0.797 |
| CSx | 0.2 | 0.694 | 0.091 | 0.703 | 0.039 | 0.807 |
| CSx | 0.3 | 0.632 | 0.099 | 0.637 | 0.057 | 0.714 |
| MuSiC | 0.02 | 0.793 | 0.084 | 0.803 | 0.016 | 0.921 |
| MuSiC | 0.05 | 0.799 | 0.083 | 0.805 | 0.018 | 0.908 |
| MuSiC | 0.1 | 0.739 | 0.089 | 0.745 | 0.032 | 0.841 |
| MuSiC | 0.2 | 0.669 | 0.095 | 0.679 | 0.04 | 0.8 |
| MuSiC | 0.3 | 0.665 | 0.101 | 0.687 | 0.028 | 0.861 |
| Scaden | 0.02 | 0.934 | 0.041 | 0.944 | 0.033 | 0.837 |
| Scaden | 0.05 | 0.925 | 0.044 | 0.936 | 0.035 | 0.825 |
| Scaden | 0.1 | 0.902 | 0.046 | 0.915 | 0.042 | 0.792 |
| Scaden | 0.2 | 0.846 | 0.054 | 0.859 | 0.051 | 0.747 |
| Scaden | 0.3 | 0.798 | 0.063 | 0.816 | 0.063 | 0.686 |

**Supplementary Table S10:** Deconvolution performance on real PBMC RNA-seq datasets PBMC1 and PBMC2. Scaden was trained on a mixture of *in silico* and real bulk RNA-seq data, the remaining tools used either scRNA-seq datasets as reference (CPM, MuSiC, CSx) or a in-built reference (CS).

| Method | Dataset | RMSE | Correlation | Slope | Intercept | CCC |
| --- | --- | --- | --- | --- | --- | --- |
| CPM | PBMC1 | 0.18 | -0.003 | -0.003 | 0.167 | -0.003 |
| CPM | PBMC2 | 0.114 | -0.203 | -0.094 | 0.182 | -0.155 |
| CS | PBMC1 | 0.147 | 0.437 | 0.491 | 0.085 | 0.434 |
| CS | PBMC2 | 0.101 | 0.594 | 0.754 | 0.041 | 0.577 |
| CSx | PBMC1 | 0.16 | 0.603 | 0.925 | 0.012 | 0.552 |
| CSx | PBMC2 | 0.13 | 0.456 | 0.67 | 0.055 | 0.424 |
| MuSiC | PBMC1 | 0.316 | -0.235 | -0.468 | 0.245 | -0.189 |
| MuSiC | PBMC2 | 0.299 | -0.197 | -0.542 | 0.257 | -0.127 |
| Scaden | PBMC1 | 0.104 | 0.722 | 0.805 | 0.032 | 0.717 |
| Scaden | PBMC2 | 0.052 | 0.855 | 0.848 | 0.025 | 0.855 |

**Supplementary Table S11:** Deconvolution performance on real PBMC RNA-seq data for Scaden models trained only on scRNA-seq simulated tissues (Scaden_SC) or on a mix of simulated and real tissue (Scaden_all).

| Method | Dataset | Celltype | RMSE | Corr. | Slope | Intercept | CCC |
| --- | --- | --- | --- | --- | --- | --- | --- |
| Scaden_SC | PBMC1 | Total | 0.131 | 0.564 | 0.644 | 0.059 | 0.559 |
| Scaden_SC | PBMC2 | Total | 0.077 | 0.684 | 0.689 | 0.052 | 0.684 |
| Scaden_all | PBMC1 | Total | 0.104 | 0.722 | 0.805 | 0.032 | 0.717 |
| Scaden_all | PBMC2 | Total | 0.052 | 0.855 | 0.848 | 0.025 | 0.855 |
| Scaden_SC | PBMC1 | Bcells | 0.033 | 0.648 | 0.172 | 0.006 | 0.083 |
| Scaden_SC | PBMC1 | CD4Tcells | 0.228 | 0.633 | 0.492 | -0.055 | 0.149 |
| Scaden_SC | PBMC1 | CD8Tcells | 0.101 | 0.603 | 0.761 | 0.108 | 0.562 |
| Scaden_SC | PBMC1 | Monocytes | 0.178 | 0.556 | 0.885 | 0.173 | 0.186 |
| Scaden_SC | PBMC1 | NK | 0.087 | 0.81 | 0.531 | 0.137 | 0.312 |
| Scaden_SC | PBMC1 | Unknown | 0.029 | 0.577 | 0.361 | 0.009 | 0.287 |
| Scaden_SC | PBMC2 | Bcells | 0.012 | 0.936 | 0.977 | 0.002 | 0.935 |
| Scaden_SC | PBMC2 | CD4Tcells | 0.145 | 0.767 | 0.682 | -0.057 | 0.119 |
| Scaden_SC | PBMC2 | CD8Tcells | 0.049 | 0.67 | 0.403 | 0.129 | 0.587 |
| Scaden_SC | PBMC2 | Monocytes | 0.078 | 0.865 | 0.994 | 0.071 | 0.558 |
| Scaden_SC | PBMC2 | NK | 0.071 | 0.629 | 0.314 | 0.14 | 0.276 |
| Scaden_SC | PBMC2 | Unknown | 0.025 | 0.247 | 0.217 | 0.044 | 0.209 |
| Scaden_all | PBMC1 | Bcells | 0.031 | 0.668 | 0.188 | 0.007 | 0.1 |
| Scaden_all | PBMC1 | CD4Tcells | 0.151 | 0.638 | 0.652 | -0.017 | 0.345 |
| Scaden_all | PBMC1 | CD8Tcells | 0.096 | 0.6 | 0.704 | 0.123 | 0.569 |
| Scaden_all | PBMC1 | Monocytes | 0.172 | 0.518 | 0.777 | 0.184 | 0.177 |
| Scaden_all | PBMC1 | NK | 0.036 | 0.804 | 0.488 | 0.058 | 0.71 |
| Scaden_all | PBMC1 | Unknown | 0.026 | 0.64 | 0.41 | 0.01 | 0.365 |
| Scaden_all | PBMC2 | Bcells | 0.013 | 0.936 | 0.94 | 0.0 | 0.917 |
| Scaden_all | PBMC2 | CD4Tcells | 0.074 | 0.772 | 0.769 | -0.005 | 0.373 |
| Scaden_all | PBMC2 | CD8Tcells | 0.051 | 0.672 | 0.398 | 0.106 | 0.562 |
| **Scaden_all** | PBMC2 | Monocytes | 0.072 | 0.895 | 1.058 | 0.049 | 0.614 |
| **Scaden_all** | PBMC2 | NK | 0.045 | 0.69 | 0.301 | 0.103 | 0.467 |
| **Scaden_all** | PBMC2 | Unknown | 0.023 | 0.241 | 0.178 | 0.043 | 0.203 |

**Supplementary Figure S6:**
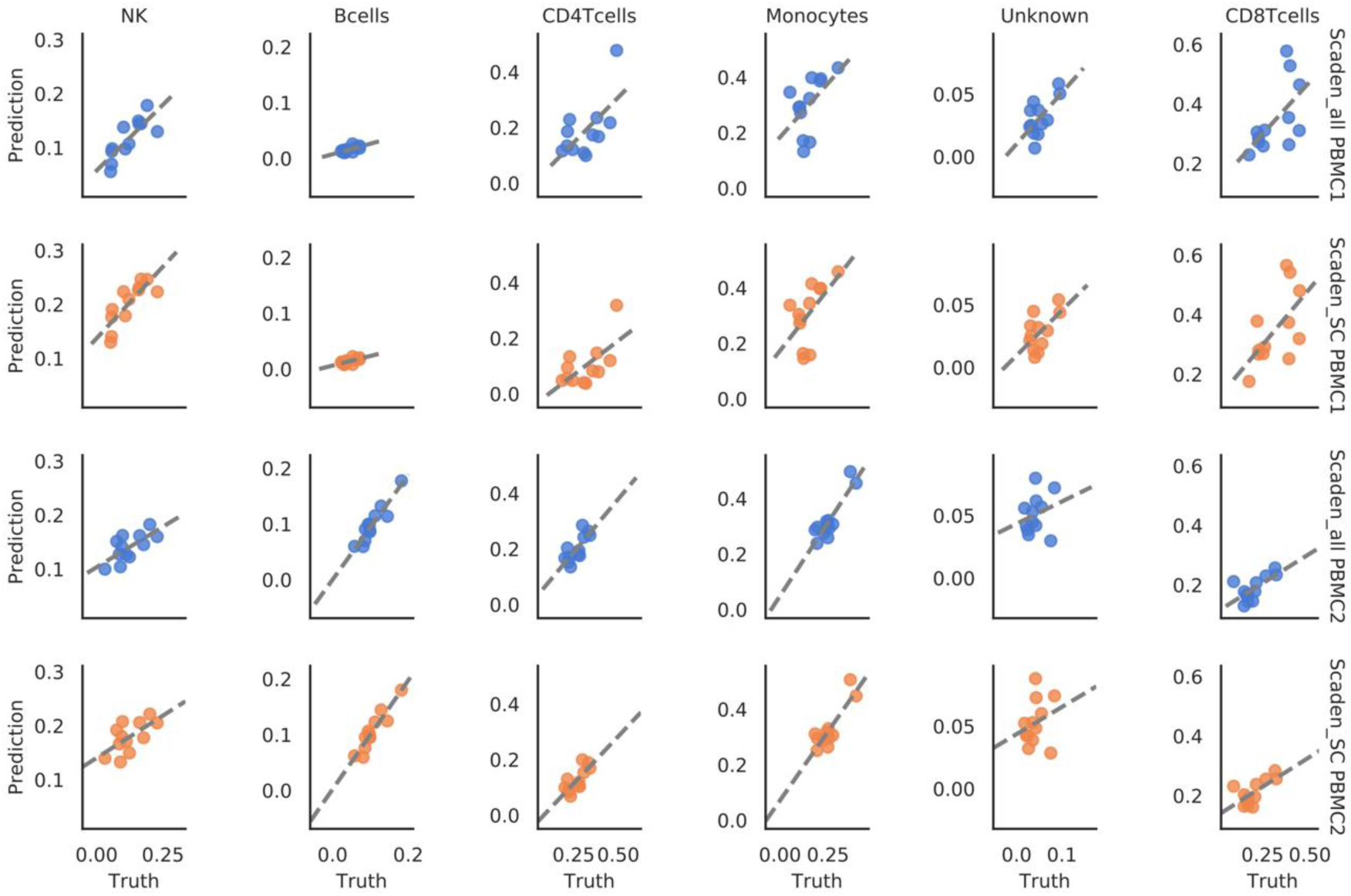
Comparison of Scaden deconvolution results on PBMC1 and PBMC2 datasets with and without (Scaden_all, Scaden_SC, respectively) bulk RNA-seq samples included in training data. Results are given on a per-cell type basis, predicted values are shown on the y-axis, ground truth is given on the x-axis.

**Supplementary Table S12:** Deconvolution performance on bulk RNA-seq data from post-mortem human brain tissue (ROSMAP). Metrics are reported for deconvolution with different reference datasets (Darmanis and Lake) and additionally with both reference datasets for Scaden (Darmanis and Lake).

| Method | Reference | CCC | RMSE | Correlation | Slope | Intercept |
| --- | --- | --- | --- | --- | --- | --- |
| Scaden | Darmanis | 0.922 | 0.056 | 0.924 | 0.992 | 0.028 |
| Scaden | Lake | 0.76 | 0.144 | 0.918 | 1.730 | 0.103 |
| Scaden | Darmanis& Lake | 0.898 | 0.064 | 0.899 | 0.948 | 0.029 |
| MuSiC | Darmanis | 0.873 | 0.089 | 0.951 | 1.445 | 0.075 |
| MuSiC | Lake | 0.65 | 0.119 | 0.652 | 0.696 | 0.078 |
| CSx | Darmanis | 0.805 | 0.123 | 0.94 | 1.663 | 0.094 |
| CSx | Lake | 0.706 | 0.183 | 0.95 | 2.130 | 0.115 |

**Supplementary Figure S7:**
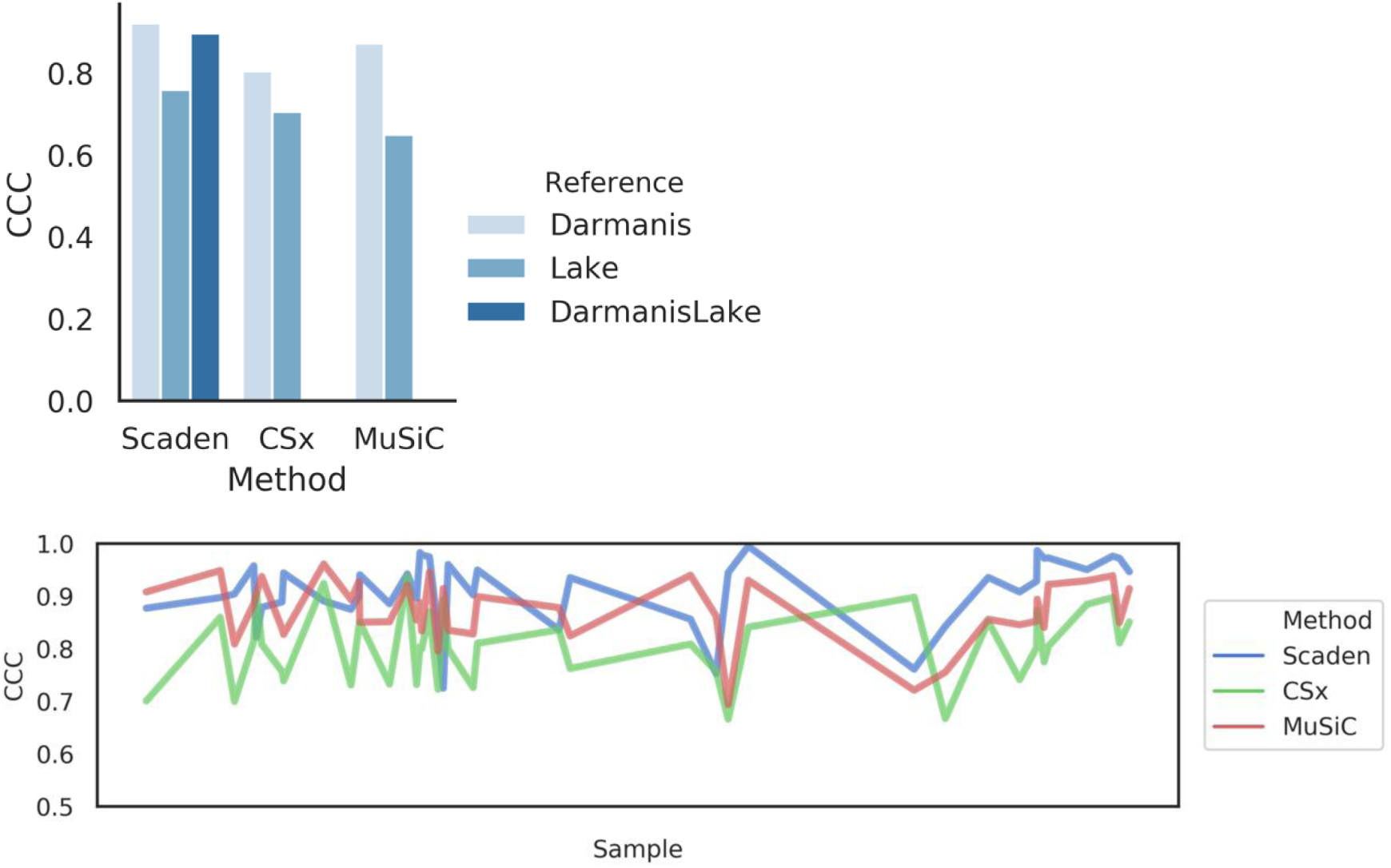
Top panel: CCC values for prediction of post-mortem human brain (ROSMAP) cell fractions for Scaden, CSx, and MuSiC on the reference data sets Darmanis (lightblue, leftmost bars) and Lake (medium-shaded blue), as well as both (for Scaden only, dark blue). Note that the Lake dataset contains only neurons, such that we fit only a subset of cell types (excitatory and inhibitory neurons, respectively). Interestingly, addition of the Lake dataset into training data affected Scaden performance only slightly. Bottom panel: Per sample results for prediction of post-morten human brain (ROSMAP) cell fractions on the Darmanis reference datasets using Scaden, CSx, and MuSiC.

**Supplementary Table S13:** Deconvolution performance comparison of CS (LM22) and Scaden on the GSE65133 Microarray dataset. Please note that the LM22 GEP used for CS was created using PBMC microarray data, while Scaden was trained on simulated scRNA-seq PBMC datasets.

| Method | Celltype | CCC | Correlation | Intercept | R2 | RMSE | Slope |
| --- | --- | --- | --- | --- | --- | --- | --- |
| CIBERSORT | Average | 0.391 | 0.612 | 0.087 | 0.396 | 0.103 | 0.516 |
| CIBERSORT | Bcells | 0.122 | 0.33 | 0.029 | 0.109 | 0.068 | 0.109 |
| CIBERSORT | CD4Tcells | 0.629 | 0.658 | 0.199 | 0.433 | 0.095 | 0.537 |
| CIBERSORT | CD8Tcells | 0.285 | 0.635 | 0.018 | 0.404 | 0.12 | 0.375 |
| CIBERSORT | Monocytes | 0.295 | 0.741 | 0.19 | 0.548 | 0.17 | 0.779 |
| CIBERSORT | NK | 0.623 | 0.698 | -0.003 | 0.487 | 0.059 | 0.78 |
| CIBERSORT | Total | 0.717 | 0.728 | 0.026 | 0.53 | 0.11 | 0.869 |
| Scaden | Average | 0.498 | 0.726 | -0.032 | 0.536 | 0.115 | 1.006 |
| Scaden | Bcells | 0.431 | 0.728 | 0.012 | 0.53 | 0.055 | 0.388 |
| Scaden | CD4Tcells | 0.64 | 0.778 | -0.195 | 0.606 | 0.153 | 1.474 |
| Scaden | CD8Tcells | 0.474 | 0.543 | 0.02 | 0.294 | 0.104 | 0.635 |
| Scaden | Monocytes | 0.43 | 0.838 | 0.033 | 0.702 | 0.191 | 1.764 |
| Scaden | NK | 0.516 | 0.741 | -0.029 | 0.549 | 0.074 | 0.77 |
| Scaden | Total | 0.705 | 0.749 | -0.015 | 0.561 | 0.126 | 1.067 |

**Supplementary Figure S8:**
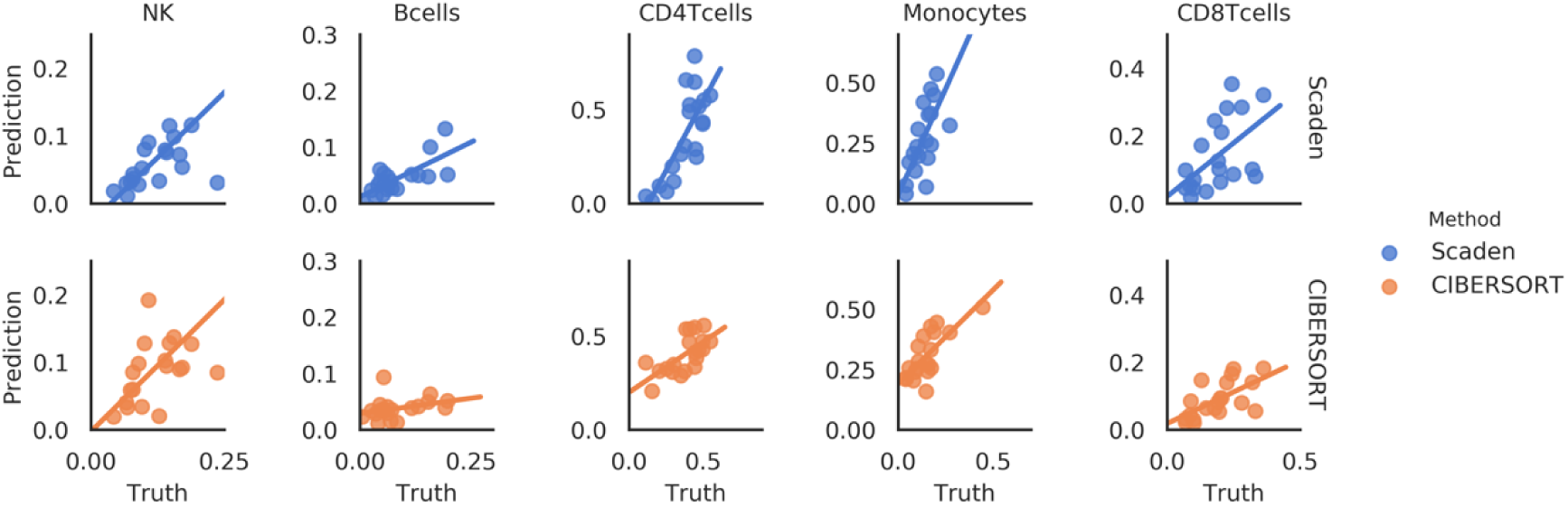
Scatterplots showing the deconvolution performance of CS (LM22) and Scaden on the GSE65133 Microarray dataset. Please note that the LM22 GEP used for CS was created using PBMC microarray data, while Scaden was trained on simulated scRNA-seq PBMC datasets.

**Supplementary Table S14:**
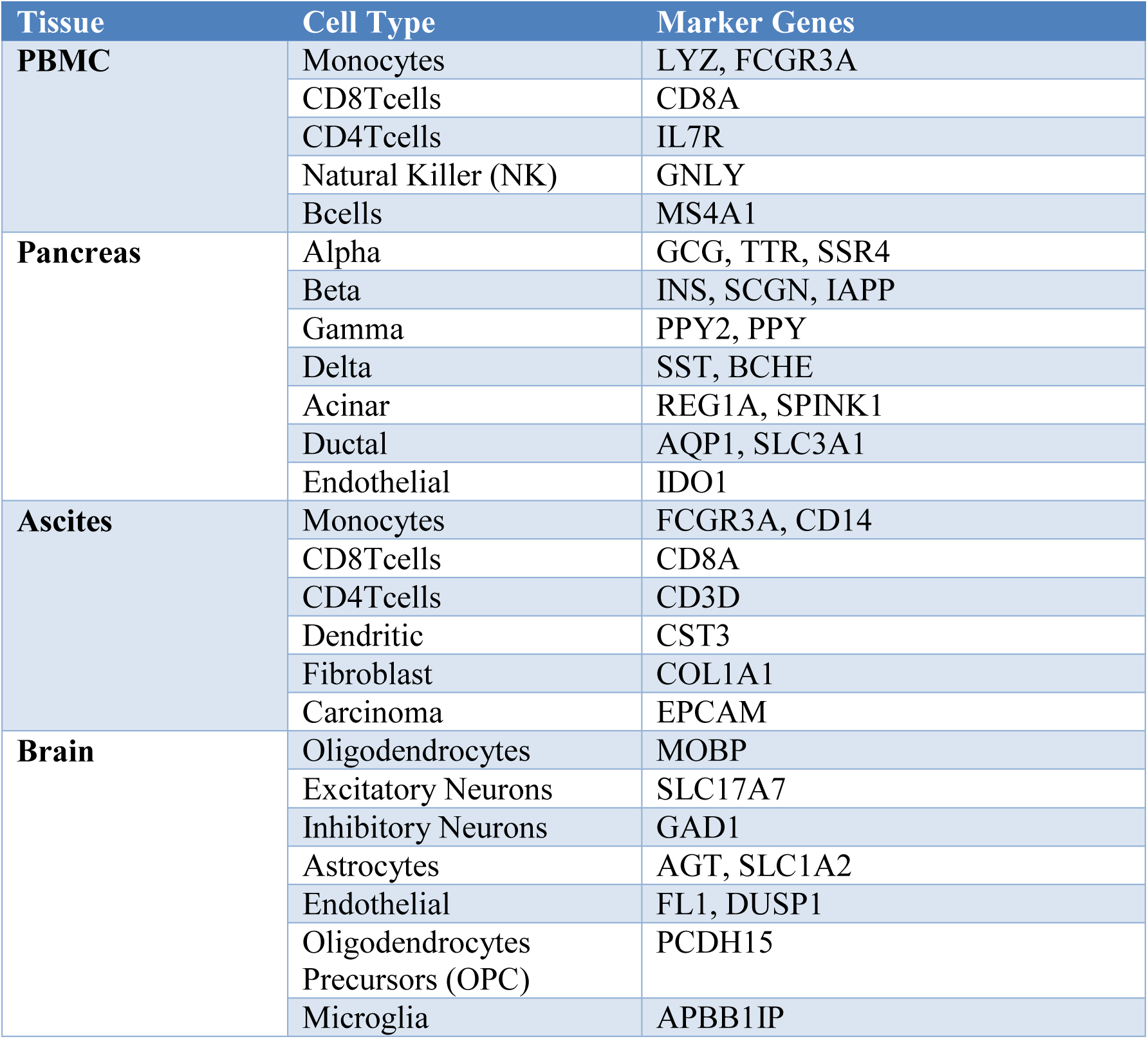
Marker genes used to define cell type populations

| Tissue | Cell Type | Marker Genes |
| --- | --- | --- |
| **PBMC** | Monocytes | LYZ, FCGR3A |
|  | CD8Tcells | CD8A |
|  | CD4Tcells | IL7R |
|  | Natural Killer (NK) | GNLY |
|  | Bcells | MS4A1 |
| **Pancreas** | Alpha | GCG, TTR, SSR4 |
|  | Beta | INS, SCGN, IAPP |
|  | Gamma | PPY2, PPY |
|  | Delta | SST, BCHE |
|  | Acinar | REG1A, SPINK1 |
|  | Ductal | AQP1, SLC3A1 |
|  | Endothelial | IDO1 |
| **Ascites** | Monocytes | FCGR3A, CD14 |
|  | CD8Tcells | CD8A |
|  | CD4Tcells | CD3D |
|  | Dendritic | CST3 |
|  | Fibroblast | COL1A1 |
|  | Carcinoma | EPCAM |
| **Brain** | Oligodendrocytes | MOBP |
|  | Excitatory Neurons | SLC17A7 |
|  | Inhibitory Neurons | GAD1 |
|  | Astrocytes | AGT, SLC1A2 |
|  | Endothelial | FL1, DUSP1 |
|  | Oligodendrocytes Precursors (OPC) | PCDH15 |
|  | Microglia | APBB1IP |

**Supplementary Table S15:** Software packages and versions used.

| Software | Version |
| --- | --- |
| **pandas** | 0.23.4 |
| **Python** | 3.6.8 |
| **Tensorflow** | 1.10.0 |
| **matplotlib** | 2.2.3 |
| **nb_conda** | 2.2.1 |
| **numpy** | 1.15.0 |
| **scipy** | 1.1.0 |
| **seaborn** | 0.9.0 |
| **anndata** | 0.6.9 |
| **scanpy** | 1.2.2 |
| **scikit-learn** | 0.20.0 |
| **ipython** | 6.5.0 |
| **python-igraph** | 0.7.1.post6 |
| **louvain** | 0.6.1 |
| **tqdm** | 4.7.2 |
| **igraph** | 0.7.1 |

**Supplementary Table S16:** Mapping of the LM22 GEP to cell types.

| Target Cell Type | LM22 Cell Types |
| --- | --- |
| B cells | B cells naive, B cells memory |
| CD8 T cells | T cells CD8, T cells follicular helper, T cells gamma delta |
| CD4 T cells | T cells CD4 naive, T cells regulatory (Tregs), T cells CD4 memory resting, T cells CD4 memory activated |
| NK | NK cells resting, NK cells activated |
| Monocytes | Monocytes, Macrophages M0, Macrophages M1, Macrophages M2 |
| Unknown | Mast cells resting, Mast cells activated, Eosinophils, T cells follicular helper, T cells gamma delta, Plasma cells, Neutrophils, Dendritic cells resting, Dendritic cells activated |

